# Taxol exploits molecular switches within tubulin to stabilize microtubules

**DOI:** 10.64898/2026.05.17.725690

**Authors:** Nicholas Vangos, Patrick DeLear, Ezekiel C. Thomas, Kristen J. Verhey, Morgan E. DeSantis, Marija Zanic, David Sept, Michael A. Cianfrocco

**Author notes:** To whom correspondence should be addressed: M.A.C.

## Abstract

Microtubules are dynamic filaments of tubulin heterodimers that comprise an essential part of the eukaryotic cytoskeleton^1^. The nucleotide state of tubulin controls microtubule dynamics: stable GTP-microtubules favor polymerization, whereas unstable GDP-microtubules drive depolymerization^2^. Anticancer compounds such as Taxol (paclitaxel) target microtubule dynamicity by preventing microtubule depolymerization^3,4^. Despite decades of work, the molecular basis of microtubule dynamics remains poorly defined. Using cryo-EM, we determined ∼2.2 Å structures of human microtubules in GTP-like (GMPCPP) and GDP states. Comparison of these two states revealed switch-like structural changes as tubulins transition from the pre-hydrolysis (GMPCPP) to the post-hydrolysis (GDP) state. Additional structure determination of Taxol-bound microtubules at ∼2.2 Å showed that Taxol binding converts the microtubule lattice into a pre-hydrolysis state by reversing the structural switches flipped during GTP hydrolysis. Focusing our analysis on the microtubule seam shows that the pre-hydrolysis conformation of GMPCPP or Taxol-GDP exhibits favorable lateral interactions at the seam, with lattice deformations clearly visible at the GDP seam. Together, our data show the existence of structural switches in tubulin that are coupled to the nucleotide state and are exploited by Taxol to stabilize microtubules into a pre-hydrolysis-like state. (191 words)

## Main

Microtubules are eukaryotic cytoskeletal filaments that provide structural rigidity and facilitate essential cellular processes, including mitosis and cargo trafficking. Heterodimers of α- and β-tubulin readily polymerize into microtubules in the presence of GTP, which is rapidly hydrolyzed to GDP upon lattice incorporation.^1^ The mature, GDP-bound microtubule lattice is intrinsically unstable and favors depolymerization. Actively growing microtubules are capped by nascent GTP-bound segments, which protect them from depolymerization. The nucleotide-driven stability of microtubules gives rise to persistent cycles of microtubule growth and shrinkage, a process known as dynamic instability.^2^ Dynamic instability is a fundamental feature of microtubules that enables their diverse set of cellular functions.

Understanding the structural basis of dynamic instability is especially important given that numerous anti-cancer compounds operate by modulating microtubule dynamics.^4^ Indeed, one of the most widely used chemotherapeutic drugs is Taxol (generic name: paclitaxel), which inhibits cell division by blocking microtubule depolymerization.^3^ Structural studies have shown that Taxol binds to a luminal site on β-tubulin near a lateral intermolecular contact.^5–7^

Microtubules assume distinct structural states when bound to different nucleotides, small molecules, or microtubule-associated proteins (MAPs).^8^ Previous cryo-EM studies have shown that microtubules assume an expanded lattice structure when bound to the non-hydrolyzable GTP analog, GMPCPP, and a compacted lattice structure when bound to GDP.^6,9,10^ Some stabilizing drugs, like Taxol, favor an expanded lattice, while others, like peloruside A, favor a compacted lattice.^7,11^ MAPs have also been observed to drive both expansion and compaction.^12–17^ Finally, recent work showed that microtubules *in situ* can be in an expanded state despite being bound to GDP in the exchangeable site.^18^

Despite their importance, the molecular mechanisms by which Taxol and nucleotide identity control microtubule stability remain poorly understood. This is due to the limited resolution (∼3 Å) of existing microtubule structures and differences in structure-determination methods. For instance, many previous structural studies utilized microtubules saturated with different MAPs. While the MAPs served to aid structure alignment during data processing, their presence on the microtubule inherently hinders interpretation, as the MAPs themselves induce structural changes in the lattice. The most common MAPs employed in these studies are kinesin motor domain (which expands the lattice^12,19^) and DCX (which compacts the lattice^14^).

This has resulted in a set of conflicting models for how GMPCPP or Taxol stabilize microtubules. Some models suggest that longitudinal changes along^6,7,9^ or lateral changes between^20–23^ protofilaments give rise to the increased stability of GMPCPP or Taxol-bound microtubules. Additionally, low resolution structures of the microtubule seam suggest that there may be nucleotide-dependent changes^7,9,24^, but without higher-resolution structural data the role of the seam in microtubule stability remains unclear.

To understand how Taxol and GMPCPP stabilize microtubules, we sought to determine the structures of human microtubules in the absence of additional factors. To this end, we developed a novel cryo-EM data analysis pipeline to determine single protofilament structures of GMPCPP-bound, GDP-bound, and Taxol-GDP-bound human microtubules at 2.2-2.3 Å resolution. These structures, which are the highest-resolution microtubule structures to date, allowed us to visualize >400 structured water molecules across two dimers, providing new insights into side-chain- and water-mediated structural changes.

Comparison of the pre-hydrolysis (GMPCPP) and post-hydrolysis (GDP) structures revealed three switch-like regions that undergo concerted structural changes to facilitate lattice compaction. Surprisingly, Taxol induces a tubulin structure that closely mimics the GMPCPP state in striking detail, reversing the switch positions in the GDP state. We also demonstrate that the GDP state exhibits staggered lattice compaction at the heterotypic microtubule seam, resulting in unfavorable lateral kinking. GMPCPP prevents these compaction-driven changes altogether, while Taxol reverses them via switch-driven expansion. Our data support a seam-centric view of microtubule stability: Taxol “flips” molecular switches to push tubulin into an expanded, pre-hydrolysis-like state, thereby stabilizing the microtubule seam and suppressing hydrolysis-driven microtubule destabilization at non-favorable heterotypic contacts. Our data show that Taxol’s anti-cancer properties stem from its ability to mimic a GTP-like state, which promotes microtubule growth while suppressing microtubule dynamics.

### GTP hydrolysis drives concerted changes in “switch” regions of tubulin

Previous microtubule structures were of limited resolution, likely due to the use of brain-derived tubulin, which is known to be a mix of tubulin isotypes with varying degrees of post-translational modifications, and whole-microtubule reconstructions, which restrict angular sampling.^6,25,26^ Without high-resolution details of specific side chain conformations and structured water molecules, the structural changes of tubulin that occur during GTP hydrolysis and Taxol binding remain unknown. To mitigate these potential issues, we 1) purified human tubulin from HeLa cells, which provides a tubulin sample with minimal post-translational modifications and isotype variety^27,28^, and 2) developed a new cryo-EM analysis pipeline. Our analysis pipeline relaxes constraints on neighboring particles, focusing instead on determining the structure of one or two protofilaments within the microtubule lattice (**Extended Data Fig. 1A**; see **Methods**). Using this new approach, we determined 2.2-2.3 Å reconstructions of human tubulin in pre-hydrolysis (GMPCPP) and post-hydrolysis (GDP) states (**Fig. 1A, Extended Data Fig. 1C&D**), enabling unprecedented insight into key structural features of the microtubule lattice and allowing us to identify more than 400 structured water molecules within each structure.

**Fig. 1.**
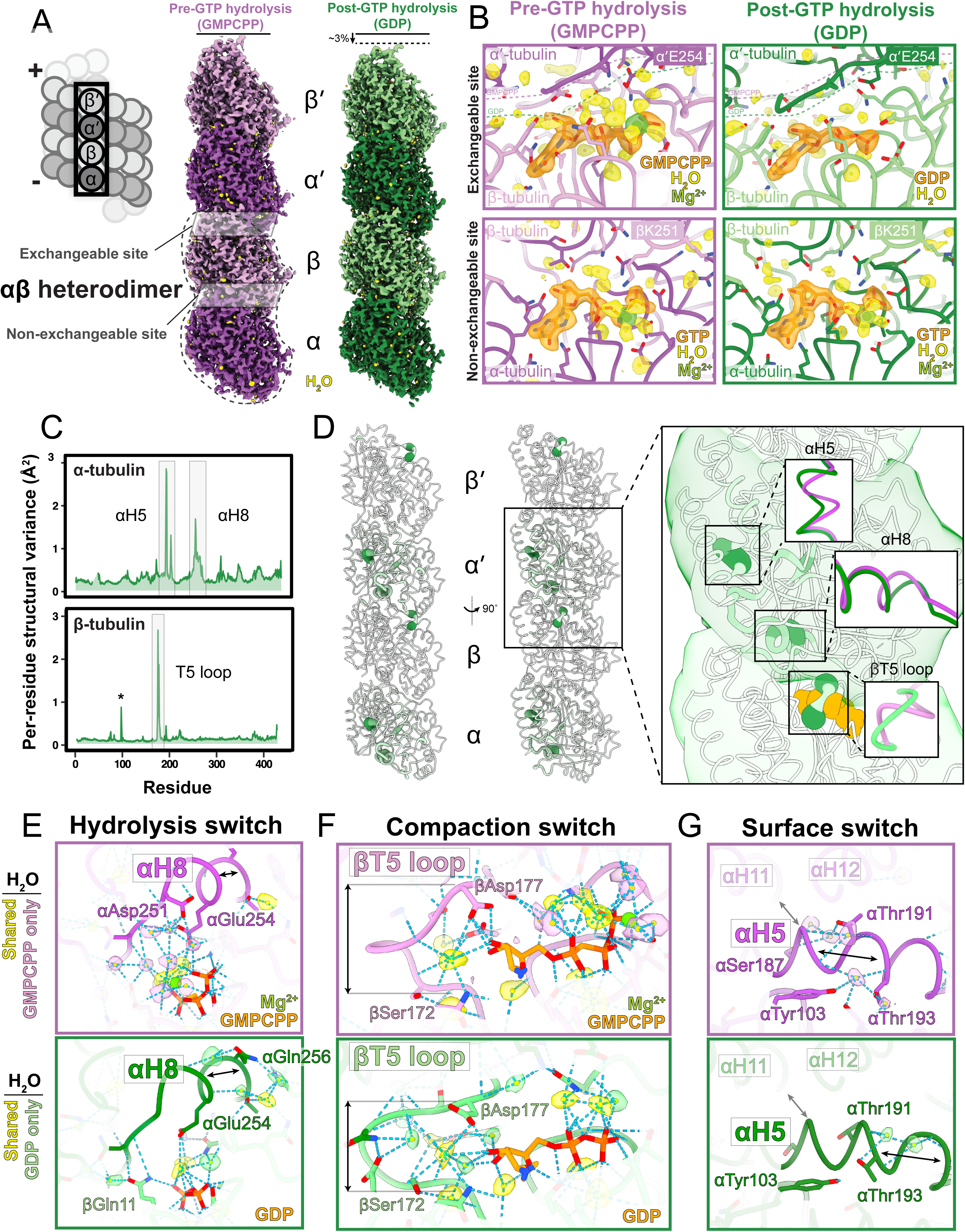
GTP hydrolysis drives concerted changes in “switch” regions via water network reorganization. (A) (Left) Schematic of microtubule structure, indicating a single protofilament comprised of two αβ heterodimers. Dashed line indicates longitudinal compaction. (Right) Cryo-EM reconstructions of single protofilaments from GMPCPP and GDP datasets at 2.2 Å and 2.3 Å, respectively. Nucleotides are orange, water molecules are yellow, and magnesium ions are lime green. (B) View of exchangeable and non-exchangeable nucleotide sites in GMPCPP and GDP single protofilament structures. Dashed lines indicate longitudinal compaction. (C) Mean per-residue structural variance comparison between GMPCPP and GDP structures for α-and β-tubulin. Boxes mark switch regions, asterisk marks nucleotide-sensitive T3 loop. (D) (left) Depiction of mean per-residue structural variance results mapped onto a single protofilament, where increased thickness and darker color indicate a greater difference between GMPCPP and GDP structures. (Right) Zoom in on the interdimer interface showing insets of switch regions within α- and β-tubulin between GMPCPP (purple) and GDP (green) structures. Nucleotide is orange. (E-G) Zoom in on switches in GMPCPP and GDP structures. Water molecules present in both structures are colored yellow, where GMPCPP or GDP-specific water molecules are colored purple and green, respectively. Sky blue dashed lines depict calculated hydrogen bonds. Black arrows depict notable structural changes.

Comparing the GMPCPP and GDP structures highlighted global and local structural changes within and between tubulin heterodimers. As previously described^6,9^, the GDP protofilament structure exhibited a ∼3% global decrease in longitudinal dimer spacing relative to GMPCPP (80.5 Å vs. 83.2 Å) (**Fig. 1A, Extended Data Fig. 1C&D**). At the exchangeable site on β-tubulin (**Fig. 1B, top left and right**), we observe clear density for GMPCPP and GDP, with the catalytic magnesium ion present only in the GMPCPP structure (**Fig. 1B, top left**). The magnesium ion bound to GMPCPP adopts an octahedral coordination geometry, coordinating four water molecules and two oxygen atoms from the β- and γ-phosphates (**Fig. 1B, top left**). At the non-exchangeable sites on α-tubulin, we observe that the magnesium coordination for GTP is nearly identical to that observed for magnesium bound to GMPCPP (**Fig. 1B, bottom left and right**), providing further validation of our cryo-EM structures and modeling.

In the exchangeable site, we observed a striking difference in the number of structured water molecules between each structure (**Fig. 1B, top left and right**). The GMPCPP exchangeable site shows 15 water molecules in its solvated network compared to 3 in the GDP structure, likely due to a loss of water molecules upon magnesium release during GTP hydrolysis. Additionally, the α-tubulin of the adjacent heterodimer in the GDP state sterically clashes with many of the structured water molecules in the GMPCPP state, suggesting that the compacted GDP lattice is incompatible with the structured waters surrounding GMPCPP (**Fig. 1B**). Interestingly, the non-exchangeable sites across the structures contain 8-9 waters, which is more than at the GDP exchangeable site but fewer than the GMPCPP exchangeable site. Together, our data suggest that the loss of γ-phosphate from the exchangeable site GTP leads to the concomitant loss of magnesium and associated water molecules, allowing the α-tubulin of the adjacent heterodimer to bind more closely and form a compacted lattice.

We have thus far used a fixed reference point (e.g., β-tubulin) to highlight the overall compaction of the GDP protofilament compared to GMPCPP. While this is consistent with prior work^6,9,20^, we reasoned that using a fixed reference point could mask localized changes within discrete sites, as the majority of α-carbon (C-α) displacement occurs between dimers (**Extended Data Fig. 1F&G**). Moreover, it seemed likely that subtle yet crucial intra-subunit conformational changes must be facilitating the larger, inter-subunit differences. To compare the specific structural differences between GMPCPP and GDP states in a reference-free manner, we calculated internal distance matrices for the C-α of each residue across all other residues within a given subunit. We then calculated the variance of the differences between these distance matrices across structural states.The per-residue structural variance provides a single per-residue metric that describes structural variation across two structures without the need for a fixed reference point (**Fig. 1C, Extended Data Fig. 1H**).

Analysis of the mean per-residue structural variance between GMPCPP and GDP states revealed three distinct regions that undergo significant structural changes (**Fig. 1C**). The peak in β-tubulin at position 95 (**Fig. 1C, asterisk**) likely describes a T3 loop interaction with the terminal phosphate of GMPCPP. We discuss this loop position below alongside the Taxol-GDP structure. Mapping the per-residue structural variance onto our GDP-bound model showed that the remaining three regions are relatively close to one another and emanate from the exchangeable nucleotide site (**Fig. 1D**). Furthermore, the structural differences identified by the reference-free analysis are also visible in a more careful reference-based model overlay (**Fig. 1D, inset**). These regions correspond specifically to the T5 loop in β-tubulin and α-helices 5 and 8 in α-tubulin.

These three regions exhibit high local deformation and propagate structural changes to surrounding domains. We therefore hypothesized that the T5 loop in β-tubulin and α-helices 5 and 8 in α-tubulin act as tubulin “switches” to mediate microtubule compaction.

### Water networks facilitate tubulin switches

Inspection of the switch regions identified above revealed unique water networks that facilitate discrete changes between nucleotide states. We first inspected α-tubulin H8, as it contains the catalytic αGlu254 necessary for GTP hydrolysis^29^ (**Fig. 1E, Movie 1**). Comparing αH8 between the GMPCPP and GDP structures shows that the helix is distorted in GDP, held in a more closed conformation around the nucleotide. Notably, we observe a structured water molecule binding αGlu256 and inserting into αH8 in the GDP state, likely stabilizing the bent helix conformation (**Fig. 1E, bottom**). The coordinated loss of water molecules surrounding the exchangeable site (**Fig. 1B**) alongside the hydrolysis of GTP to GDP likely drives compaction of α-tubulin around the nucleotide by bending αH8, which is stabilized by a water molecule. Given its importance in directly facilitating hydrolysis of the nucleotide, we have named this region the “hydrolysis switch.”

Further away from the exchangeable nucleotide site, we see changes in the β-tubulin T5 loop in its interaction with the ribose ring of the nucleotide (**Fig. 1F, Movie 2**). In the GMPCPP structure, the β-tubulin T5 loop adopts a more open conformation with βAsp177 strongly bound to both ribose hydroxyl groups in an upward conformation (**Fig. 1F, top**). In the GDP state, the T5 loop adopts a more compacted structure, recruiting additional water molecules to bind and reposition the GDP ribose hydroxyls downward (**Fig. 1F, bottom**). Due to the hydrolysis-driven increase in structured water molecules within the T5 loop, there are far more hydrogen bonds present in the loop in the GDP state. Given the crucial position of this switch at the inter-dimer interface, its leverage against the adjacent α-subunit, and its open-to-closed conformational change, we have named this the “compaction switch” as it mediates the transition of the adjacent α-subunit to compact on β-tubulin.

Beyond the switches located directly adjacent to the nucleotide, we observe a third switch at a distal site within α-tubulin at αH5 (**Fig. 1D, Movie 3**). Prior to hydrolysis, the helix is open/bent between αSer187 and αThr191, stabilized by water molecules that interact with both side chains and the Cα backbone (**Fig. 1G, top**). In the GDP structure, we see the loss of these waters (**Fig. 1G, bottom**). Instead, new water molecules bind between αThr193 and αGlu196, shifting the bend further down the helix. These structural changes in αH5 are adjacent to and accompanied by changes in the packing of surface helices αH11 and αH12 against tubulin. Given the position of this switch on the microtubule surface, we named this region the “surface switch.”

### Taxol exploits molecular switches to reverse microtubule compaction

We next wanted to determine how the binding of Taxol to β-tubulin affects these switch-like regions. Using the same pipeline established above, we determined the structure of Taxol bound to tubulin at 2.2 Å (**Fig. 2A, Extended Data Fig. 1E**). Inspection of the interaction between Taxol and β-tubulin at this resolution allowed unambiguous identification of Taxol’s binding pose (**Fig. 2B&C**). We observe that the two benzoyl rings within Taxol (2-benzoyl and N-benzoyl) facilitate cross-bridging interactions involving helices βH6, βH7, and βH1, ultimately sandwiching βHis227. βHis227 makes two types of interactions with Taxol: a π-π stacking with the 2-benzoyl ring and a hydrogen bond with the N-benzoyl carbonyl group (**Fig. 2B&C**). We also observe an ordered water molecule that hydrogen-bonds to the other N moiety of βHis227, located between βHis227 and βAsp224, which likely further stabilizes the conformation of βHis227. The 2-benzoyl and N-carbonyl groups are present in all taxanes used in the clinic: docetaxel, paclitaxel, and cabazitaxel, further suggesting a conserved importance with regard to the mechanism of action. On the distal ends of Taxol, we see that the benzoyl rings make hydrophobic interactions with βLeu215 on βH6 and βVal23 on βH1. Beyond interactions between Taxol’s benzoyl rings within tubulin, we see two additional interactions between Taxol and tubulin. The oxetane ring in Taxol interacts with the amide backbone of βThr274, thus allowing Taxol to interact both with βH6 and βH8, creating an interface between two different structural elements. On the other side of Taxol, the carbonyl of βArg359 binds to the 2’-hydroxyl within the C13 ester side chain. The high occupancy and density quality allows unambiguous interpretation of these interactions, unlike other Taxol-microtubule reconstructions.^7,23^ Together, these interactions help Taxol span multiple helices within β-tubulin while also making specific side-chain contacts.

**Fig. 2.**
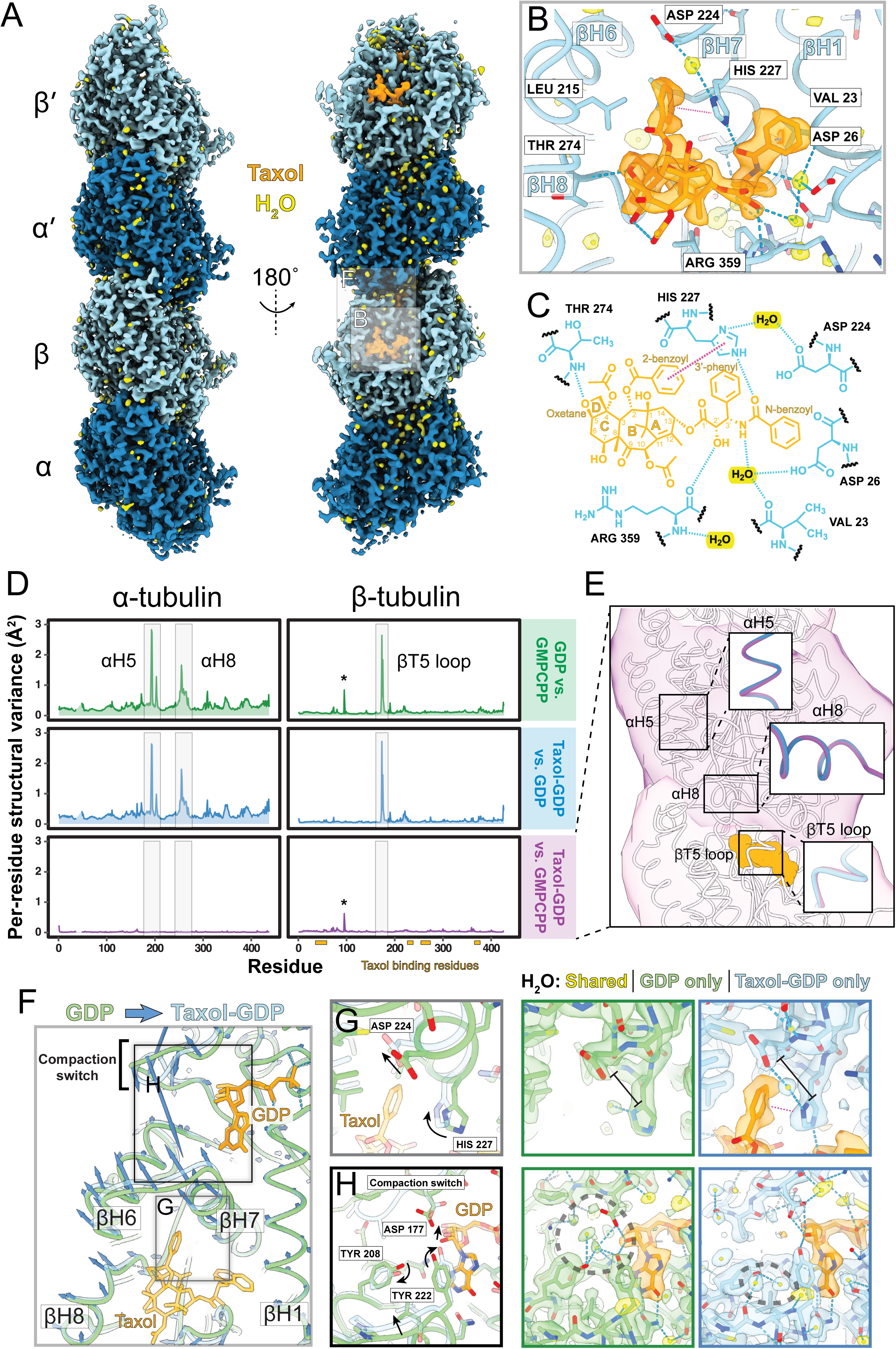
Taxol exploits molecular switches to reverse microtubule compaction. (A) Cryo-EM reconstruction of a single protofilament from Taxol-GDP dataset at 2.2 Å. (B) Binding pose of Taxol within β-tubulin. Density for Taxol (orange) and water molecules (yellow) are shown alongside the atomic model for β-tubulin. Blue dashed lines are hydrogen bonds; the magenta dashed line is π-π stacking. (C) Schematic diagram showing Taxol’s interaction with β-tubulin and structured water molecules. (D) Mean per-residue structural variance profiles for GMPCPP vs. GDP, Taxol-GDP vs. GDP, and Taxol-GDP vs. GMPCPP. Boxes mark switch regions, asterisk marks nucleotide-sensitive T3 loop. (E) Depiction of per-residue structural variance results mapped onto a single protofilament, where increased thickness and darker color indicate a greater difference between GMPCPP and GDP structures. Zoom in on the interdimer interface showing insets of switch regions within α- and β-tubulin between Taxol-GDP (blue) and GMPCPP (purple). Nucleotide is orange. Asterisk marks nucleotide-sensitive T3 loop. (F) Comparison of GDP to Taxol-GDP 1-protofilament structures near the Taxol-binding site showing C-α displacement vectors in dark blue (amplified 4-fold). (G) View of βHis227 in GDP vs. Taxol-GDP showing 2’-benzoyl group of Taxol interaction with βHis227 and the incorporation of a structured water molecule. Black arrows and lines indicate notable structural changes. (H) View of exchangeable site nucleotide ribose ring with resides at the top of βH7 and the compaction switch (top). Black arrows indicate notable structural changes. Dotted gray circles indicate notable changes in water networks.

After establishing the interaction interface between Taxol and β-tubulin, we next wanted to compare the structure of Taxol-bound tubulin with the pre- and post-hydrolysis tubulin structures. Mapping the per-residue structural variance between the Taxol-GDP, GMPCPP, and GDP structures showed a striking similarity in the GMPCPP vs. GDP and GDP vs. Taxol-GDP profiles (**Fig. 2D**). The same three switch regions identified above appear in the GDP vs. Taxol-GDP profile, suggesting a conserved mechanism of expansion. This is corroborated by consistently low per-residue structural variance scores in the Taxol-GDP vs. GMPCPP profile (average difference ∼0.2 Å) (**Fig. 2D & 2E**), with the only difference arising from an interaction between the T3 loop in β-tubulin and the γ-phosphate of GMPCPP (**Fig. 2D asterisk, Extended Data Fig. 2**). Importantly, the three switch regions across the Taxol-GDP and GMPCPP states are in the exact same position (**Fig. 2E**), suggesting that Taxol reverses these switches to facilitate lattice expansion. These results are surprising given that 1) the Taxol-GDP exchangeable site has GDP bound and 2) Taxol binds distal to the inter-tubulin interface.

Of the tubulin switches identified above, the compaction switch (β-tubulin T5 loop) is the most proximal to the Taxol binding site (**Fig. 2F**). Taxol’s extensive interaction with βHis227 repositions βH7 via a water molecule that binds between βHis227 and βAsp224, shifting βH7 to reposition βTyr222 (**Fig. 2G, Movie 4**). At the nucleotide, we see that βTyr222 repositions and stabilizes the ribose ring within the nucleotide in an ‘up’ configuration (**Fig. 2H, Movie 5**), which mirrors that of the GMPCPP structure. These changes also likely destabilize the extensive water network at the post-hydrolysis compaction switch (**Fig. 2H**, **Fig. 1F**), further driving tubulin to a pre-hydrolysis state. This shift in βH7 likely couples Taxol binding to tubulin switches and, by extension, microtubule lattice expansion.

### Taxol reverses staggered compaction and angular deformation at the microtubule seam

Next, we wanted to determine how structural changes within tubulin manifest across protofilaments to affect tubulin lattice structure. There are two kinds of lateral microtubule contacts: homotypic and heterotypic.^30^ Homotypic lattices consist of like-like contacts and make up most of the microtubule lattice, while heterotypic lattices represent the asymmetric point of contact within a 13-protofilament microtubule known as the seam.^31^ We adapted our analysis pipeline to determine structures of two adjacent protofilaments that arise from homotypic (non-seam) and heterotypic (seam) lattices from within our 13-protofilament datasets of GMPCPP, GDP, and Taxol-GDP (**Extended Data Fig. 1A**; see **Methods**).

First, we compared homotypic lattice structures for GMPCPP (2.4 Å) and GDP (2.2 Å) (**Fig. 3A**). Mapping longitudinal displacement of C-α positions from GMPCPP to GDP showed no changes for the neighboring (α’’β’’) tubulin but concerted compaction in the GDP state for both α’β’ and α’’’β’’’ tubulins (**Fig. 3A, left, Movie 6**). We did not observe notable changes in lateral interactions between these lattices. Mapping C-α displacement from GDP to Taxol-GDP (2.3 Å) showed synchronous tubulin expansion in the Taxol-GDP state without lateral changes (**Fig. 3A, right, Movie 6**), in effect reversing the hydrolysis-driven compaction. Moreover, a comparison between Taxol-GDP and GMPCPP homotypic lattices showed no discernible longitudinal differences, further supporting a shared structural state (**Extended Data Fig. 3**).

**Fig. 3.**
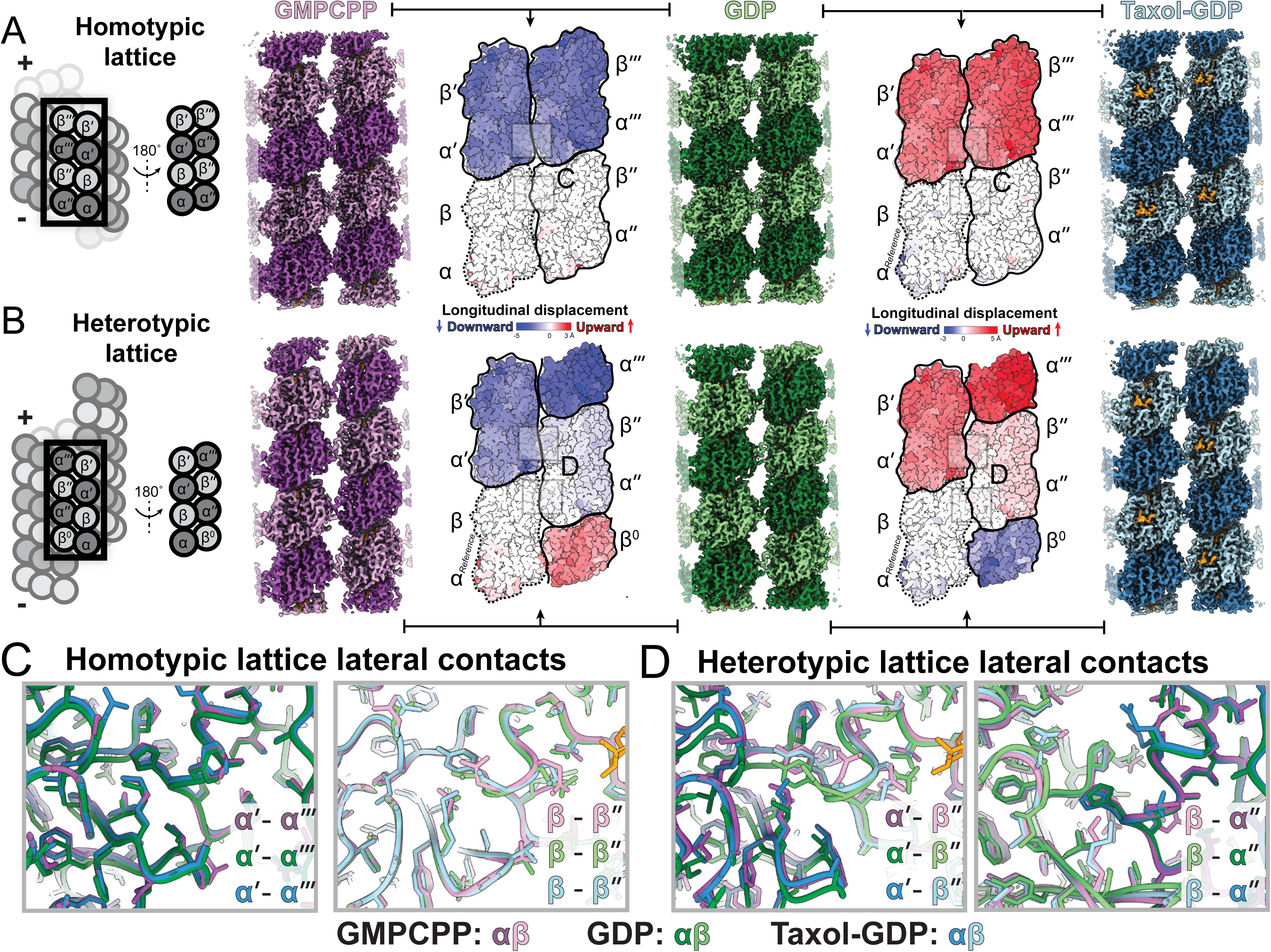
Staggered compaction at the microtubule seam perturbs α-β lateral contacts. (A) Homotypic lattices from two profilament cryo-EM structures of GMPCPP (2.4 Å), GDP (2.2 Å), and Taxol-GDP (2.3 Å). (Left) Schematic of a two-protofilament structure, indicating that structures are viewed from the lumenal side of the microtubule. (Right) Comparison between each model shows the longitudinal component of C-α displacement colored onto the surface of a two protofilament model relative to the αβ dimer. Blue is downward displacement, red is upward displacement, and white is no displacement. Black outlines indicate heterodimers. Boxes relate to (C) and (D). (B) Comparison of heterotypic lattices from GMPCPP (3.1 Å), GDP (2.7 Å), and Taxol-GDP (2.8 Å) colored in the same manner as (A). (Left) Schematic of heterotypic lattice location on a microtubule. (C & D) Zoom-ins of homotypic (C) and heterotypic (D) interprotofilament lateral lattice contacts between α and β-tubulin for regions indicated in (A) & (B). Models are aligned via the right-most subunit.

Comparison of longitudinal C-α differences for the heterotypic lattices in GMPCPP (3.2 Å), GDP (2.7 Å), and Taxol-GDP (2.8 Å) states revealed a more complex pattern of structural changes (**Fig. 3B**). Relative to the reference tubulin dimer (αβ), we see that the longitudinally adjacent tubulin heterodimer (α’β’) shows compaction comparable to the homotypic lattice. Unlike the homotypic lattice, the nearest tubulin dimer on the neighboring protofilament (α’’β’’) is offset by one subunit. As a result, the largest differences occur in the next sphere of tubulins (α’’’ and β^0^), where these tubulin heterodimers experience inter-dimer compaction. Inspection of the Taxol-GDP heterotypic lattice relative to GDP shows that Taxol-GDP exhibits staggered expansion of the seam, relieving the staggered compaction in the GDP lattice (**Fig. 3B, Movie 7**).

To explore where heterotypic vs. homotypic lattice differences arise across these three states, we compared inter-protofilament contacts (**Fig. 3C & 3D**). Superimposing α-α and β-β contacts from the homotypic lattice structures shows that the backbone and side positions of these contacts remain comparable across all states (**Fig. 3C**), consistent with the synchronous compaction/expansion across homotypic GMPCPP, GDP, and Taxol-GDP lattices (**Fig. 3A**). The largest difference between protofilaments occurs in the heterotypic lattice between α- and β-tubulin (**Fig. 3D**). With structures aligned to β-tubulin (β’’), we see that the adjacent α-tubulin (α’) in the GDP lattice is significantly displaced relative to the GMPCPP and Taxol-GDP structures (**Fig. 3D, left**). This contrasts with the heterotypic β-α-tubulin contacts, which appear consistent across all structures (**Fig. 3D, right**). These comparisons suggest that the α-β lateral contacts are uniquely compromised at the microtubule seam due to uneven strain induced by hydrolysis-driven staggered compaction in the GDP state.

Protofilament contact angles also identify the GDP seam as an outlier (**Fig. 4).** We observed only small differences in lateral contact angles across the homotypic lattice from each structural state (**Fig. 4B, Movie 8**). However, the heterotypic lattices exhibit a dramatic and striking difference in contact angle between GMPCPP and GDP states (**Fig. 4C, Movie 9**). This angular difference is unique to the heterotypic lattice, suggesting that this is a seam-specific phenomenon. We hypothesize that this lateral deformation is indicative of a weakened, less-favorable heterotypic lattice at the post-hydrolysis, GDP seam. The addition of Taxol relieves this angular strain at the seam, returning the contact angle to that of the pre-hydrolysis state. The heterotypic lattice angles in stable lattice structures - 152.8° for GMPCPP and 153.5° for Taxol-GDP - more closely reflect the homotypic lattice angles (154.0° avg.) than does the heterotypic GDP lattice (147.2°), suggesting that the post-hydrolysis heterotypic lattice represents a weakened lattice state.

**Fig. 4.**
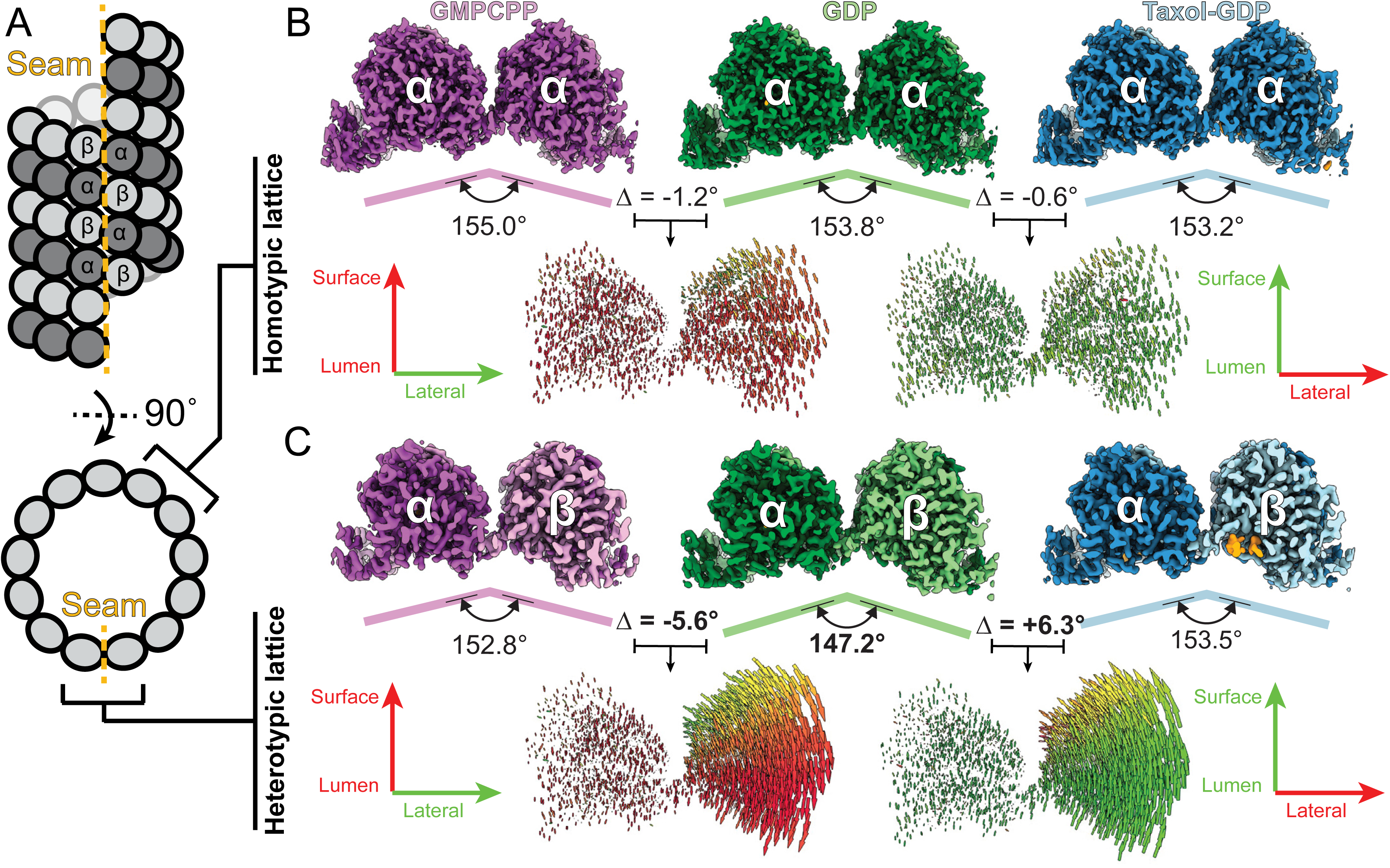
Taxol relieves hydrolysis-driven strain at the microtubule seam. (A) Schematic showing a microtubule alongside the top-down views of homotypic and heterotypic lattices. (B) Top-down views of homotypic lattice 2-profilament models for GMPCPP (purple), GDP (green), and Taxol-GDP (blue) states. (Bottom) Pairwise comparisons showing vectors between C-α positions. The vectors are dilated by a factor of distance/(max_distance*2) to emphasize larger relative changes and are colored according to the X and Y displacement with green and red, respectively. (C) Top-down view of heterotypic lattice 2-protofilament models for GMPCPP (purple), GDP (green), and Taxol-GDP (blue) show alongside vector pairwise comparisons, as in (B).

## Discussion

In this work, we used cryo-EM to determine high-resolution structures of human-derived microtubules, enabling the discovery of structural switches within tubulin that mediate microtubule transition from pre-hydrolysis to post-hydrolysis states. Additionally, we show that the anti-cancer compound Taxol exploits these switches to “flip” microtubules from a GDP state to a GTP-like state. Finally, we show that structural transitions between these states are strongly manifested at the microtubule seam, with the GTP-like and Taxol-GDP states adopting a favorable seam conformation, whereas the GDP state exhibits perturbed lateral contacts and an aberrant contact angle.

We propose a unified model of microtubule stability that connects the nucleotide state to discrete and concerted changes in molecular switches across tubulin heterodimers, facilitating the larger structural changes spanning microtubule lattices (**Fig. 5**). Our data suggest that nucleotide hydrolysis initiates these structural changes: release of GTP’s terminal phosphate and its coordinate magnesium flips the “hydrolysis” switch in α-tubulin, which propagates structural rearrangements along the tubulin lattice; the β-tubulin “compaction” switch mediates inter-molecular changes at the nucleotide binding interface, while the “surface” switch facilitates intra-molecular changes (**Fig. 5A**). Microtubule-targeting agents such as Taxol can exploit these switches to reverse nucleotide-driven compaction, likely flipping the “compaction” switch first and triggering a cascade through the “hydrolysis” and “surface” switches (**Fig. 5B**). By mimicking a GTP-like state, Taxol induces a conformation on tubulin that promotes polymerization, suppressing microtubule dynamics which are necessary for cell growth.

**Fig. 5.**
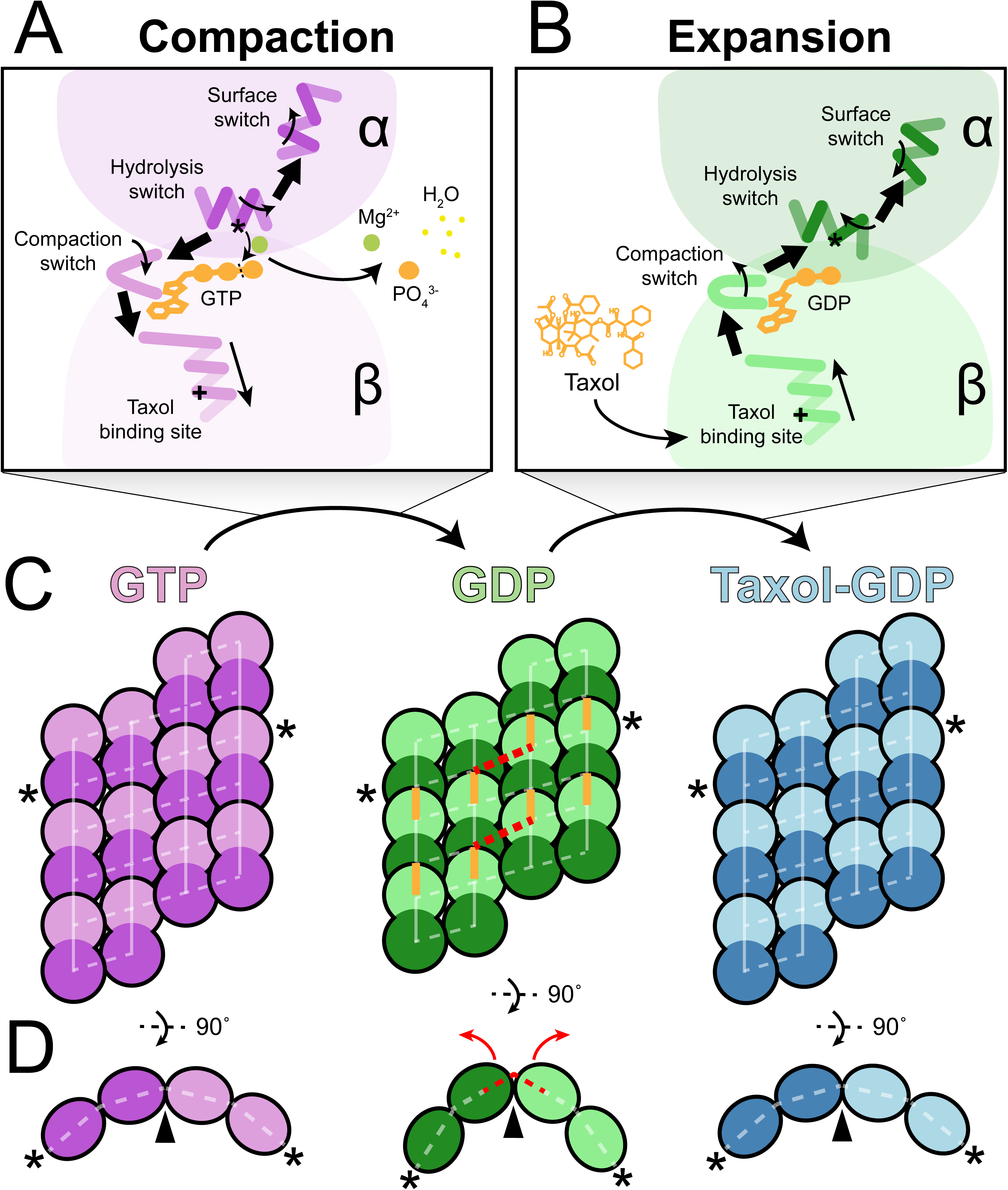
Model of microtubule structural changes induced by GTP hydrolysis and reversed by Taxol binding. (A) Model of changes to tubulin switches that occur between GMPCPP and GDP structural states during the process of microtubule compaction. Small arrows indicate local structural changes, large arrows indicate propagative changes between regions. Asterisk indicates catalytic residue Glu254. Plus sign indicates His227. (B) Reversion of tubulin switches by the binding of Taxol to convert the GDP structure to an expanded state. (C) Model showing lattice changes, highlighting the synchronous conformational changes for homotypic lattices and the asymmetric staggering at the heterotypic lattice in the GDP state. While lines indicate favorable contacts, yellow lines indicate compaction, and red dotted lines indicate non-favorable contacts. (D) Top-down view showing the weakened heterotypic lattice in the GDP structure due to staggered compaction and inter-protofilament bending. Asterisks in (C) denote the contiguous lateral strip of tubulin shown in (D).

Additionally, our analysis of 2-protofilament lattice structures strongly links microtubule lattice compaction and expansion to microtubule stability. We see that homotypic lattices undergo concerted longitudinal changes without impacting lateral contacts (**Fig. 5C**). For the first time, we observed that the heterotypic lattice at microtubule seams undergoes staggered compaction upon GTP hydrolysis (**Fig. 5C**), perturbing α-β lateral contacts. This results in a weakened lattice structure, evidenced by a sharper contact angle between protofilaments in the unstable post-hydrolysis GDP seam lattice (**Fig. 5D**). The unique lattice at the GDP seam could recruit specific proteins, such as SPEF1 in cilia.^32^ Our findings suggest that microtubule instability arises from strain at the microtubule seam and that Taxol stabilizes microtubules by relieving this strain.

Our structure of Taxol bound to human tubulin maps the taxane interaction site in molecular detail and shows that structured water molecules are critical to the interaction (**Fig. 2B&C**), highlighting the need for high-resolution, ligand-bound structures. The proposed model expands our understanding of taxane-tubulin interactions and provides the foundation for future medicinal chemistry approaches to improve the chemotherapeutic efficacy of taxanes.

Furthermore, we show that Taxol unambiguously induces a GTP-like state within the microtubule lattice. Taxol-GDP microtubules mimic both longitudinal and lateral conformations of GMPCPP (**Extended Data Fig. 3**). Taxol’s ability to induce this structural state within tubulin leads to a tubulin conformation that promotes microtubule growth without shrinkage. These data suggest that other lattice-expanding stabilizers, such as the clinically-relevant drug ixabepilone (brand name Ixempra), may stabilize microtubules in a similar manner to induce apoptosis. Non-expanding stabilizers like peloruside A and laulimalide, which bind a distinct site near lateral microtubule contacts, might stabilize microtubules by directly strengthening lateral interactions, thereby alleviating lateral strain at the seam.

We believe our work introduces a new framework for understanding the connections among nucleotide state, lattice conformation, and microtubule stability. Future work that utilizes our approach of human tubulin with high-resolution (< 2.5 Å) structures will connect the mechanism of GTP hydrolysis with microtubule-associated proteins and to map changes in the microtubule lattice. Finally, our analysis framework will facilitate the determination of microtubule structure in cells, enabling future work that connects microtubule lattice states to *in situ* cellular states.

## Acknowledgements

We thank Cianfrocco, DeSantis, R. Ohi, Sept, and Verhey labs for critical discussions of this work in addition to the U-M cryo-EM facility staff. We also thank Chi-Min Ho, Elizabeth Kellogg, and Gary Brouhard for feedback on this work. The research reported in this publication was supported by the University of Michigan Cryo-EM Facility (U-M Cryo-EM). U-M Cryo-EM is grateful for support from the U-M Life Sciences Institute and the U-M Biosciences Initiative. We acknowledge the following funding sources: S10OD030275 (N.V. & M.A.C), R01GM141119 (N.V. & M.A.C.), T32GM145470 (N.V.), R35GM131744 (K.J.V.), PF-24-1320851-01-CCB (E.C.T.), R01GM136822 (P.D & D.S.), R35GM119552 (M.Z.), and R35GM146739 (M.E.D.).

## Author contributions

N.V. prepared, imaged, and collected cryo-EM data. N.V. performed single particle analysis and built atomic models. E.C.T. purified HeLa tubulin. P.D. and D.S. developed and performed structural analysis. N.V., M.E.D., K.J.V., M.Z., D.S., and M.A.C. wrote the paper with input from all authors. M.A.C. supervised the study and conceived of the project.

## Methods

### GST-TOG protein expression and purification

To purify tubulin from HeLa cells, we adapted the approach of using a TOG affinity column ^28,33^. pGEX-6p-1Stu2(1-590)TOG1/2 was expressed in 2L of BL21 DE3 cells. Briefly, these were grown at 37°C for ∼3 hours, induced at OD 0.6 with 0.4 mM IPTG, and cooled to 20°C overnight. Cell pellets were harvested and resuspended in 50mL of lysis buffer (1X PBS, 1mM MgCl_2_, 0.5% Triton-X100, 1mM ATP, 1mM DTT, 0.2mM PMSF, pH 8.0). Resuspension was sonicated in a cooled steel cup at 60% amplitude for 18 cycles of 10s on / 20s off (3 min total lysis time). Lysate was centrifuged at 39,000 rcf for 30 min at 4°C. Supernatant was filtered with a 1.0 µm syringe filter and batch bound to 5mL of equilibrated glutathione resin for 1.5 hours. The incubation mixture was then flowed through a packable PD-10 column. Settled resin was washed with 25mL of Triton-free lysis buffer (1X PBS, 1mM MgCl_2_, 1mM ATP, 1mM DTT, 0.2mM PMSF, pH 8.0) and 25mL of wash buffer (1X PBS, 1mM MgCl_2_, 0.2mM PMSF, pH 8.0) before eluting with 3 x 5mL of elution buffer (1X PBS, 5mM reduced glutathione, pH 8.0). Eluent was dialyzed against 2L of dialysis buffer (100mM NaHCO_3_, 100mM NaCl, pH 8.2) in preparation for column coupling. Dialyzed sample was concentrated to >5mg/mL with an Amicon 10kDa cutoff filter, flash frozen, and stored at -80°C.

### Generation of GST-TOG column

GST-TOG protein aliquots totaling >25mg total (more is better) were thawed and diluted to ∼5mg/mL in dialysis buffer. Protein was coupled to ∼1mL of NHS-activated Sepharose 4 Fast Flow resin following manufacturer instructions. Briefly, MgCl_2_-6H_2_O powder was added to the pooled GST-TOG protein to a final concentration of 80mM, and the mixture was gently inverted to dissolve powder. ∼1mL of resin was washed with 10mL of cold 1mM HCl. GST-TOG solution was then flowed and cycled back over the resin for 30-40 min at 27°C. 3mL of quenching solution (0.5M ethanolamine, 0.5M NaCl, pH 8.3) was added to stop the coupling reaction, and the column was washed with 10mL of 6X PBS and 10mL of 1X of PBS.

### HeLa tubulin purification via GST-TOG column

HeLa S3 cells were grown in suspension culture as described.^27,28^ 800mL of cells yielding 2-3mL cell pellets were processed at a time. The cell pellet was resuspended in 20mL of 1X BRB80 (80mM PIPES, 1mM EGTA, 1mM MgCl_2_, pH 6.8) + 0.1mM GTP + 1mM DTT + 1X Roche protease inhibitor cocktail + 500 units of benzonase. Resuspension was sonicated at 20% amplitude for 3 x 10s on / 20s off (30s total lysis time). Lysate was centrifuged at 165,000 rcf for 40 min at 4°C. Supernatant was filtered with a 1.0 µm syringe filter and loaded onto the GST-TOG column equilibrated with 25mL of 1X BRB80. Flow through was reapplied to the column twice to ensure exhaustive tubulin binding. Column was then sequentially washed with 20mL of wash buffer 1 (1X BRB80, 0.1mM GTP), 20mL of wash buffer 2 (1X BRB80, 0.01mM GTP), and 10mL of wash buffer 3 (1X BRB80, 0.1mM GTP, 5mM ATP). After a 10-minute incubation with wash buffer 3, the column was washed with 10mL of wash buffer 2. The column was then eluted with 2 x 2.5mL elution buffer (1X BRB80, 0.01mM GTP, 500mM (NH_4_)_2_SO_4_). Eluent was immediately desalted to 1X BRB80 using PD-10 desalting columns to minimize tubulin exposure to high salt, concentrated to >5mg/mL using an Amicon 10kDa filter, diluted with glycerol to 10% final concentration, and snap frozen in 10μL aliquots. The GST-TOG column was regenerated by successive washes with 25mL of 1X PBS, 25mL of 10X PBS, and 25mL of 1X PBS. The column can be stored in 1X PBS + 50% glycerol at -20°C for 2+ years.

### GDP microtubule sample preparation

1x 10μL aliquot of ∼5mg/mL HeLa tubulin in 1X BRB80 + 10% glycerol was thawed rapidly by hand and put on ice. 1μL of 10mM GTP was then added to the tubulin, tapping the tube to mix, and incubated on ice for 5 min. The polymerization mixture was then transferred to a 37°C heat block and allowed to incubate for 45 min. After polymerization, the microtubule mixture was spun at 17,000 rcf at 25°C for 22 min. This spin was done in a centrifuge adjacent to the vitrobot to minimize time between spin completion, pellet resuspension, and sample vitrification. Supernatant was carefully pipetted off, and the pelleted microtubules were gently resuspended in 10μL of warm 1X BRB80 + 25% glycerol + 2mM GTP using a p10 pipette with the tip cut ∼1mm from the end to minimize shearing. Pellet was resuspended for ∼1 min, and 4μL of resuspension was plunged on a Quantifoil 1.2/1.3 400 Au grid using a Thermo Fisher Vitrobot Mark IV with the following plunging parameters: 60s wait time, 5s blot time, 20 blot force, 37°C, 100% humidity.

### Taxol-GDP sample preparation

1x 10μL aliquot of 5mg/mL HeLa tubulin in 1X BRB80 + 10% glycerol was thawed rapidly by hand and put on ice. 1μL of 10mM GTP was then added to the tubulin, tapping the tube to mix, and incubated on ice for 5 min. The polymerization mixture was then transferred to a 37°C heat block and allowed to incubate for 15 min. 1μL of 1μM Taxol in DMSO was then added and mixed, and the solution was allowed to incubate for 10 min. 1μL of 10μM Taxol in DMSO was then added and mixed, followed by another 10 min incubation. 1μL of 100μM Taxol in DMSO was then added and mixed before a final 15 min incubation. After polymerization, the microtubule mixture was spun at 17,000 rcf at 25°C for 22 min. Supernatant was carefully pipetted off, and the pelleted microtubules were gently resuspended in 20μL of warm 1X BRB80 + 25% glycerol + 10μM Taxol using a p20 pipette with the tip cut ∼1mm from the end to minimize shearing. Pellet was resuspended for ∼1 min, and 4μL of resuspension was plunged on a Protochip 1.2/1.3 C-Flat 300 Au grid using a Thermo Fisher Vitrobot Mark IV with the following plunging parameters: 60s wait time, 2s blot time, 0 blot force, 30°C, 90% humidity.

### GMPCPP sample preparation

1x 10μL aliquot of 5mg/mL HeLa tubulin in 1X BRB80 + 10% glycerol was thawed rapidly by hand and put on ice. The following mixture was then prepared on ice: 10μL HeLa tubulin, 0.5μL 100mM MgCl_2_, 0.5μL 100mM DTT, 4μL 10mM GMPCPP, and 35μL 1X BRB80. The polymerization mixture was incubated on ice for 5 min before transfer to a 37°C heat block and incubation for 45 min. After polymerization, the microtubule mixture was spun at 17,000 rcf at 25°C for 22 min. After the spin, the supernatant was carefully pipetted off, and the pelleted microtubules were gently resuspended in 20μL of warm 1X BRB80 + 25% glycerol using a p10 pipette with the tip cut ∼1mm from the end to minimize shearing. Pellet was resuspended for ∼1 min, and 4μL of resuspension was plunged on a Quantifoil 2/1 200 Cu grid using a Thermo Fisher Vitrobot Mark IV with the following plunging parameters: 0s wait time, 4s blot time, 20 blot force, 37°C, 90% humidity.

### Cryo-EM data collection

The samples were imaged using a FEI Titan Krios (300 kV) equipped with a K3 detector and energy filter (20 eV slit size) (Gatan) using automated data collection SerialEM^34^. All movies were acquired at 105,000 X magnification (0.834 Å/pixel, no objective aperture, ∼1.9 sec exposure, 50 frames, ∼50 e/Å^2^, -0.8 to -2.0 μm defocus range). 4,282 movies were collected for the GDP dataset, 3,197 movies were collected for the Taxol-GDP dataset, and 12,822 movies were collected across two GMPCPP datasets (6,270 and 6,552). All datasets were preprocessed on-the-fly utilizing motion-correction and CTF estimation in WARP^35^ to confirm desired defocus range during data collection. The GDP dataset collection exhibited periodic misfocusing during collection and was thus filtered by defocus (under -2.5 μm) and estimated resolution (better than 6Å) in WARP, resulting in 2,879 usable movies. Datasets were then imported into RELION-5.0, where all following steps were performed unless mentioned otherwise.

### Cryo-EM whole-microtubule data processing

All datasets were treated similarly leading up to whole-microtubule helical refinement (**Extended Data Fig. 4**). Imported movies were motion corrected using RELION’s own implementation of MotionCor2^36^. CTFFIND-4.1.14^37^ was used to estimate the CTF of each micrograph. The Taxol-GDP and GMPCPP datasets were filtered to better than 6Å maximum resolution, yielding 3,004 micrographs for Taxol-GDP and 12,034 micrographs for both GMPCPP datasets (5,593 and 6,441). To generate good 2D templates for particle picking, a 3D reference of a 13-protofilament microtubule was used to template pick on a random subset of 100 micrographs. This resulted in 23,879 particles for GDP, 41,863 particles for Taxol-GDP, and 14,521 particles for both GMPCPP datasets (11,985 and 2,536). These initial particles were extracted at 4x binning with a 180 pixel (600Å) box size and subjected to 2D classification. 4-14 good classes were used as 2D templates to re-pick on their entire respective datasets. This resulted in 126,763 particles for GDP, 159,284 particles for Taxol-GDP, and 508,886 particles for both GMPCPP datasets (397,419 and 111,467). We found that dataset-derived 2D classes worked much better as picking templates than imported 2D classes from other datasets. Full-dataset particle picks were extracted at 4x binning to make runtimes more manageable prior to resolution-dependent steps downstream. GDP particles were then subjected to an optional round of 2D classification to clean the particle stack, yielding 104,935 particles. However, it is advised to skip this step if you want more accurate distributions of protofilament numbers, as manual curation introduces bias.

Microtubule particles for each dataset were subjected to a supervised 3D classification using 6 synthetic references to sort by protofilament number, as previously described ^26^. In agreement with the literature, we saw mostly 13-protofilament particles for the GDP and Taxol-GDP datasets, while the GMPCPP datasets were mostly 14-protofilament. It is advisable to use the first-iteration classes from this job to obtain more accurate protofilament distributions, as subsequent iterations prioritize only the best and most abundant particles. However, the final-iteration class for GDP 13-protofilament microtubules was used instead, as it yielded a high quality reconstruction. This step resulted in 49,269 13-protofilament particles for GDP, 100,778 13-protofilament particles for Taxol-GDP, and 45,267 13-protofilament particles for both GMPCPP datasets (36,035 and 9,232).

Each particle stack was subjected to a round of 2D classification to remove poor-quality particles, resulting in 43,190 particles for GDP, 71,948 particles for Taxol-GDP, and 42,403 particles for both GMPCPP datasets (33,464 and 8,939). The cleaned 13-protofilament particles were then re-extracted unbinned with a 720 pixel (600 Å) box size and subjected to 3D refinement with applied helical symmetry (search range: -27.4° to -28.0°, 8.9 Å to 9.7 Å). This yielded a 3.45Å GDP reconstruction, a 3.68 Å Taxol-GDP reconstruction, and 3.96 Å & 4.38 Å GMPCPP reconstructions. Particles were then CTF Refined (per-micrograph defocus, per-micrograph astigmatism) and subjected to Bayesian polishing (default parameters). Subsequent 3D refinement with applied helical symmetry yielded a 3.28 Å GDP reconstruction, a 3.28Å Taxol-GDP reconstruction, and 3.70 Å & 4.17 Å GMPCPP reconstructions. Particles were then symmetry expanded by 26-fold to account for all possible subunit positions using the helical parameters from the final refinement jobs (**Table 1**), yielding 1,122,940 particles for GDP, 1,870,648 particles for Taxol-GDP, and 1,102,478 particles across both GMPCPP datasets (870,064 and 232,414). Symmetry expanded particles were subjected to signal subtraction with re-centering on a 400 pixel (∼334 Å) box using either a 1- or 2-protofilament mask. Subtracted stacks were combined for the GMPCPP datasets. This resulted in 2 signal subtracted particle stacks for each experimental condition. All subsequent 3D refinements were performed without applied symmetry.

**Table 1.** Cryo-EM data collection, processing, and refinement statistics.

|  | GDP HeLa MT |  |  | Taxol-GDP HeLa MT |  |  | GMPCPP HeLa MT |  |  |
| --- | --- | --- | --- | --- | --- | --- | --- | --- | --- |
| Data collection and processing | 1-pf | 2-pf<br>(homotypic) | 2-pf<br>(heterotypic) | 1-pf | 2-pf<br>(homotypic) | 2-pf<br>(heterotypic) | 1-pf | 2-pf<br>(homotypic) | 2-pf<br>(heterotypic) |
| PDB ID | XXXX | XXXX | XXXX | XXXX | XXXX | XXXX | XXXX | XXXX | XXXX |
| EMDB ID | XXXXX | XXXXX | XXXXX | XXXXX | XXXXX | XXXXX | XXXXX | XXXXX | XXXXX |
| Microscope | Titan Krios G3 |  |  |  |  |  |  |  |  |
| Voltage (kV) | 300 |  |  |  |  |  |  |  |  |
| Detector | K3 Summit |  |  |  |  |  |  |  |  |
| Magnification | 105,000 |  |  |  |  |  |  |  |  |
| Electron exposure (e<sup>-</sup>/Å<sup>2</sup>) | ~50 |  |  |  |  |  |  |  |  |
| Frame number | 50 |  |  |  |  |  |  |  |  |
| Exposure rate (e<sup>-</sup>/pixel/s) | ~12 |  |  |  |  |  |  |  |  |
| Calibrated pixel size (Å) | 0.834 |  |  |  |  |  |  |  |  |
| Defocus range (µm) | 0.7-2.2 |  |  | 0.5-2.0 |  |  | 0.5-2.0 |  |  |
| Symmetry imposed | C1 |  |  | C1 |  |  | C1 |  |  |
| Movies | 2,879 |  |  | 3,197 |  |  | 12,711 |  |  |
| Initial particle images (before symmetry-expansion) | 126,763 |  |  | 159,284 |  |  | 508,886 |  |  |
| Final particle images | 629,845 | 351,622 | 89,998 | 719,914 | 616,487 | 99,070 | 313,939 | 481,602 | 37,672 |
| Map resolution (Å) | 2.42 | 2.42 | 2.85 | 2.42 | 2.47 | 3.01 | 2.49 | 2.59 | 3.44 |
| FSC threshold | 0.143 | 0.143 | 0.143 | 0.143 | 0.143 | 0.143 | 0.143 | 0.143 | 0.143 |
| Map sharpening factor (Å<sup>2</sup>) | -50.3 | -44.4 | -46.6 | -53.2 | -52.1 | -56.6 | -47.1 | -54.1 | -60.0 |
| Density-modified map resolution (Å) | 2.19 | 2.14 | 2.65 | 2.23 | 2.26 | 2.77 | 2.29 | 2.35 | 3.14 |
| Density-modified FSC threshold | 0.5 | 0.5 | 0.5 | 0.5 | 0.5 | 0.5 | 0.5 | 0.5 | 0.5 |
| Helical symmetry (13-pf MT map) |  |  |  |  |  |  |  |  |  |
| Single subunit rise (Å) | 9.29 |  |  | 9.61 |  |  | 9.60 |  |  |
| Helical twist (°) | -27.71 |  |  | -27.68 |  |  | -27.68 |  |  |
| Dimer rise (Å) | 80.46 |  |  | 83.23 |  |  | 83.08 |  |  |
| Axial twist (°) | -0.16 |  |  | 0.12 |  |  | 0.10 |  |  |
| Refinement |  |  |  |  |  |  |  |  |  |
| Initial models (AlphaFold ID) |  |  |  |  |  |  |  |  |  |
| α-tubulin | Human_tubA1B_AF-P68363-F1-model_v4 |  |  |  |  |  |  |  |  |
| β-tubulin | Human_tubB4B_AF-P68371-F1-model_v4 |  |  |  |  |  |  |  |  |
| Model resolution (Å) | 2.01 | 2.07 | 2.52 | 2.06 | 2.2 | 2.65 | 2.08 | 2.24 | 3.08 |
| FSC threshold | 0.5 | 0.5 | 0.5 | 0.5 | 0.5 | 0.5 | 0.5 | 0.5 | 0.5 |
| Model composition |  |  |  |  |  |  |  |  |  |
| Non-hydrogen atoms | 13,582 | 27,000 | 26,984 | 13,790 | 27,332 | 27,316 | 13,597 | 27,088 | 76,056 |
| Protein residues | 1,702 | 3,404 | 3,404 | 1,708 | 3,416 | 3,416 | 1,700 | 3,412 | 3,412 |
| Waters | 428 | 16 | 0 | 426 | 16 | 0 | 403 | 32 | 0 |
| Ligands | 6 | 12 | 12 | 7 | 14 | 14 | 8 | 16 | 16 |
| B factors (Å<sup>2</sup>) |  |  |  |  |  |  |  |  |  |
| Proteins | 11.51 | 10.83 | 27.28 | 9.51 | 13.68 | 52.00 | 4.80 | 11.83 | 74.56 |
| Waters | 11.17 | 2.04 | - | 10.44 | 1.36 | - | 5.36 | 2.96 | - |
| Ligands | 3.40 | 2.11 | 15.53 | 11.00 | 12.90 | 55.95 | 1.83 | 3.27 | 61.17 |
| R.M.S. deviations |  |  |  |  |  |  |  |  |  |
| Bond lengths (Å) (# > 4 ) | 0.005 (2) | 0.005 (4) | 0.005 (4) | 0.006 (10) | 0.006 (20) | 0.007 (21) | 0.005 (6) | 0.005 (14) | 0.005 (13) |
| Bond angles (°) (# > 4 ) | 1.133 (3) | 1.105 (3) | 1.133 (12) | 1.169 (7) | 1.180 (11) | 1.180 (14) | 1.090 (1) | 1.075 (6) | 0.1083 (6) |
| Validation |  |  |  |  |  |  |  |  |  |
| MolProbity score | 1.65 | 1.85 | 1.93 | 1.53 | 1.82 | 2.06 | 1.50 | 1.66 | 2.26 |
| Clashscore | 7.71 | 9.06 | 9.36 | 7.01 | 10.16 | 10.04 | 7.45 | 8.55 | 11.58 |
| Poor rotamers (%) | 1.98 | 2.18 | 3.76 | 1.52 | 2.34 | 4.43 | 1.31 | 1.82 | 4.50 |
| Ramachandran plot |  |  |  |  |  |  |  |  |  |
| Favored (%) | 98.52 | 97.42 | 98.07 | 99.00 | 97.94 | 97.79 | 98.81 | 98.47 | 96.93 |
| Allowed (%) | 1.42 | 2.55 | 1.84 | 1.00 | 2.06 | 2.21 | 1.19 | 1.53 | 3.07 |
| Disallowed (%) | 0.06 | 0.03 | 0.09 | 0.00 | 0.00 | 0.00 | 0.00 | 0.00 | 0.00 |
| Rama-Z score | 0.48 | 0.88 | 0.43 | 0.76 | -0.02 | 0.43 | 0.48 | 0.25 | 0.21 |

### Cryo-EM 1-protofilament data processing

All datasets were treated similarly during 1-protofilament refinement (**Extended Data Fig. 5, 6, & 7, left**).1-protofilament subtracted particles were backprojected to generate reference maps for subsequent refinements. Initial 3D refinement yielded 2.72 Å GDP, 2.76 Å Taxol-GDP, and 2.83 Å GMPCPP maps with a blurred α-tubulin and β-tubulin register. To sort the tubulin register, 2 high-resolution (6 Å) reference maps were generated using the molmap command in ChimeraX^38^ by fitting human tubulin (pdb: 5IJ0) into the initial mixed-register map in both possible lattice configurations. Refined particles were then subjected to a supervised 3D classification without alignment, using the previous 2 molmaps as unfiltered 6 Å references and a regularisation parameter T of 200 (100 for GMPCPP). This step leverages high-resolution prior information, which is the reason for the high T value and unfiltered references. The GMPCPP particles were subjected to an additional round of supervised 3D classification due to an initial class fraction >>50% (66.4%). The resulting classes are representative of the 2 different lattice configurations. Particles in class 1 (box centered on inter-dimer interaction) for each condition were 3D refined to 2.74 Å for GDP, 2.73 Å for Taxol-GDP, and 2.83 Å for GMPCPP.

Optics groups were enumerated for each image shift position using a custom python script, and particles were CTF refined using per-particle defocus, per-micrograph astigmatism, beamtilt estimation, trefoil estimation, and 4th order aberration estimation. CTF refined particles were then 3D refined to 2.42 Å for GDP, 2.42 Å for Taxol-GDP, and 2.49 Å for GMPCPP. Final halfmaps were exported and sharpened using the resolve cryo-EM^39^ job in phenix^40^ using default parameters, yielding density maps of 2.19 Å for GDP, 2.23 Å for Taxol-GDP, and 2.29 Å for GMPCPP. These final maps were resampled by a factor of 1.5 in Coot.^41^ Structure validation shown in **Extended Data Fig. 8A, 9A, & 10A**.

### Cryo-EM 2-protofilament homotypic lattice data processing

GDP and Taxol-GDP datasets were treated similarly during 2-protofilament homotypic refinement (**Extended Data Fig. 5 & 6, middle**). 2-protofilament subtracted particles were backprojected to generate reference maps for subsequent refinements. Initial 3D refinement yielded 2.67 Å GDP, 2.76 Å Taxol-GDP, and 2.83 Å GMPCPP maps with a blurred α-tubulin and β-tubulin register. To sort the tubulin register, 4 high-resolution (6 Å) reference maps were generated using the molmap command in ChimeraX^38^ by fitting human tubulin into the initial mixed-register map in all possible lattice configurations. Refined particles from GDP and Taxol-GDP conditions were then subjected to a supervised 3D classification without alignment, using the previous 4 molmaps as unfiltered 6Å references and a regularisation parameter T of 200. The resulting classes are representative of 2 sorted homotypic lattice configurations and 2 mixed lattices. Particles in class 1 (box centered on inter-dimer interaction) for both conditions were 3D refined to 2.65 Å for GDP and 2.76 Å for Taxol-GDP.

The GMPCPP unsorted particles were instead subjected to iterative supervised classification without alignment (T=100) using focus masks to prioritize 1-protofilament at a time (**Extended Data Fig. 7, middle**). The left protofilament was first sorted into 2 its two possible configurations. Class 1 particles (box centered on inter-dimer interaction) were enriched for the same register in the right protofilament. 2 rounds of class 1 enrichment were then combined to and 3D refined to 2.83 Å.

Optics groups were enumerated for each image shift position using a custom python script, and particles were CTF refined using per-particle defocus, per-micrograph astigmatism, beamtilt estimation, trefoil estimation, and 4th order aberration estimation. CTF refined particles were then 3D refined to 2.42 Å for GDP, 2.47 Å for Taxol-GDP, and 2.59 Å for GMPCPP. Final halfmaps were exported and sharpened using the resolve cryo-EM^39^ job in phenix^40^ using default parameters, yielding density maps of 2.14 Å for GDP, 2.26 Å for Taxol-GDP, and 2.35 Å for GMPCPP. These final maps were resampled by a factor of 1.5 in Coot^41^. Structure validation shown in **Extended Data Fig. 8B, 9B, & 10B**.

### Cryo-EM 2-protofilament heterotypic lattice data processing

To sort the heterotypic microtubule lattice, iterative supervised classifications with focus masking were necessary in all cases. Our exact approaches changed as subsequent datasets proved more challenging, but the central idea of leveraging high-resolution subunit structures in known lattice configurations holds true. This demonstrates the flexibility of our approach, both for microtubule structures and for other structurally similar but compositionally heterogeneous repeating structures.

To determine the GDP heterotypic lattice, a 2-subunit focus mask was made for two lateral central subunits (**Extended Data Fig. 5, right**). Subtracted and refined particles with a blurred α-tubulin and β-tubulin register were subjected to supervised 3D classification without alignment, using the aforementioned 4 molmaps as unfiltered 6 Å references and a regularisation parameter T of 100. This was done in an effort to sort particles into all 4 possible lattice configurations: β-β, β-α, α-α, and α-β. Particles in classes 2 and 4, corresponding to the desired heterotypic lattice states, were cleaned via 3 iterative rounds of the same job until only heterotypic lattice remained. At this point, 89,998 particles in class 2 were 3D refined to 3.01 Å, exhibiting a clear heterotypic lattice.

For the Taxol-GDP heterotypic lattice, focus masks corresponding to one of the two protofilaments were used to sort the lattice in sequence (**Extended Data Fig. 6, right**). Subtracted and refined particles with a blurred α-tubulin and β-tubulin register were subjected to supervised 3D classification without alignment, using 2 molmaps as unfiltered 6Å references and a regularisation parameter T of 100. Class 1 particles (box centered on inter-dimer interaction) were taken forward, undergoing 3 iterative rounds of supervised classification focusing on the other protofilament. Particles associated with class 2 (box centered on a heterodimer) were enriched here. 99,070 particles from the final supervised classification were 3D refined to 3.12 Å, exhibiting a clear heterotypic lattice.

For the GMPCPP heterotypic lattice, focus masks corresponding to one of the two protofilaments were used to sort the lattice in parallel (**Extended Data Fig. 7, right**). Subtracted and refined particles with a blurred α-tubulin and β-tubulin register were subjected to supervised 3D classification without alignment, using 2 molmaps from ChimeraX^38^ as unfiltered 6 Å references and a regularisation parameter T of 100. This was done for each protofilament using the same starting stack of particles, with the left protofilament undergoing 2 iterative rounds of classification. Left protofilament particles from class 1 (box centered on inter-dimer interaction) and right protofilament particles from class 2 (box centered on a heterodimer) were then inner joined using a custom Python script. The resulting particle stack only contains particles present in both input stacks. These 37,672 particles were then 3D refined to 3.55 Å, exhibiting a clear heterotypic lattice.

Optics groups were enumerated for each image shift position using a custom Python script, and particles were CTF refined using per-particle defocus, per-micrograph astigmatism, beamtilt estimation, trefoil estimation, and 4th order aberration estimation. CTF refined particles were then 3D refined to 2.85 Å for GDP, 3.01 Å for Taxol-GDP, and 3.44 Å for GMPCPP. Final halfmaps were exported and sharpened using the resolve cryo-EM^39^ job in phenix^40^ using the “density modify unsharpened maps” option, yielding density maps of 2.65 Å for GDP, 2.77 Å for Taxol-GDP, and 3.14 Å for GMPCPP. These final maps were resampled by a factor of 1.5 in Coot^41^. Structure validation shown in **Extended Data Fig. 8C, 9C, & 10C**.

### Model building and refinement

Alphafold2^42^ models for human α-tubulin 1B (Human_tubA1B_AF-P68363-F1-model_v4) and β-tubulin 4B (Human_tubB4B_AF-P68371-F1-model_v4) were used as starting models. These models were rigid-body fit into each density map using ChimeraX^38^. Models were then iteratively refined into the final density maps using real-space refinement in Phenix (version 1.21.1-5286) and Coot (version 0.9.8.95), prioritizing density fit, Ramachandran outliers, geometry, and rotamer outliers in that order. GTP, GDP, GMPCPP, and Taxol ligands were added and fit in Coot. Waters were modeled using the “Find waters” option in Coot with the default h-bond distance range (2.4-3.2 Å) and the default density threshold of 1.4 rmsd. Waters were then manually curated one by one in Coot, added or removed as needed, with the density threshold set to 2.0 rmsd. Waters were removed unless they met the following requirements: 1. there was still density at 2.0 rmsd; 2. they made at least 1 h-bond, preferably with a non-water; and 3. they survived a local real-space refinement in coot.

### Per-residue structural variance

Per-residue structural variance was calculated by taking the variance of pairwise distances for each residue from the difference-distance matrices comparing states (**Extended Data Fig. 1H**). Difference-distance matrices were calculated by measuring the pairwise distances of alpha carbons within a given chain and state (ex. Chain A1 in GDP) and subtracting the corresponding pairwise distances of the same chain in a separate state (ex. Chain A1 in GDP-Taxol). The per-residue structural variance for each residue is shown in **Fig. 1C** and **2D**. Our analysis was done in R with help from the bio3D library^43^, although this can be done in many ways.

### Axial twist and protofilament angle calculations

Axial microtubule twists were calculated from the unsharpened, whole-microtubule maps in ChimeraX using the following command: >measure symmetry #<VOLUME-spec> minimumCorrelation 0.9 helix 82,0,opt. Inter-protofilament angles for 2-protofilament structures analyzed in **Fig. 4** were calculated from rotational matrices of fitted β-tubulin subunits using the matchmaker command in ChimeraX.

### Model colorization and vectorization

.defattr, .pb, and .bild files for depicting C-α per-residue structural variance coloration and displacement vectorization were generated using a custom Python script.

## Data Availability

Coordinates for 1-protofilament, 2-protofilament homotypic, and 2-protofilament heterotypic GDP microtubule structures are available at the Protein Data Bank (PDB) with accession codes XXXX, XXXX, and XXXX, respectively. Cryo-EM density maps for 1-protofilament, 2-protofilament homotypic, and 2-protofilament heterotypic GDP microtubule structures are available at the Electron Microscopy Data Bank (EMDB) with accession codes XXXX, XXXX, and XXXX, respectively. The cryo-EM dataset for GDP microtubules is available at the Electron Microscopy Public Image Archive (EMPIAR) with accession code XXXX.

Coordinates for 1-protofilament, 2-protofilament homotypic, and 2-protofilament heterotypic Taxol-GDP microtubule structures are available at the PDB with accession codes XXXX, XXXX, and XXXX, respectively. Cryo-EM density maps for 1-protofilament, 2-protofilament homotypic, and 2-protofilament heterotypic Taxol-GDP microtubule structures are available at the EMDB with accession codes XXXX, XXXX, and XXXX, respectively. The cryo-EM dataset for Taxol-GDP microtubules is available at the EMPIAR with accession code XXXX.

Coordinates for 1-protofilament, 2-protofilament homotypic, and 2-protofilament heterotypic GMPCPP microtubule structures are available at the PDB with accession codes XXXX, XXXX, and XXXX, respectively. Cryo-EM density maps for 1-protofilament, 2-protofilament homotypic, and 2-protofilament heterotypic GMPCPP microtubule structures are available at the EMDB with accession codes XXXX, XXXX, and XXXX, respectively. The cryo-EM dataset for GMPCPP microtubules is available at the EMPIAR with accession code XXXX.

## Code Availability

Detailed cryo-EM processing pipeline and scripts for cryo-EM processing and structural analysis are available at GitHub (https://github.com/cianfrocco-lab/nvangos).

## Figure legends

**Extended Data Fig. 1.**
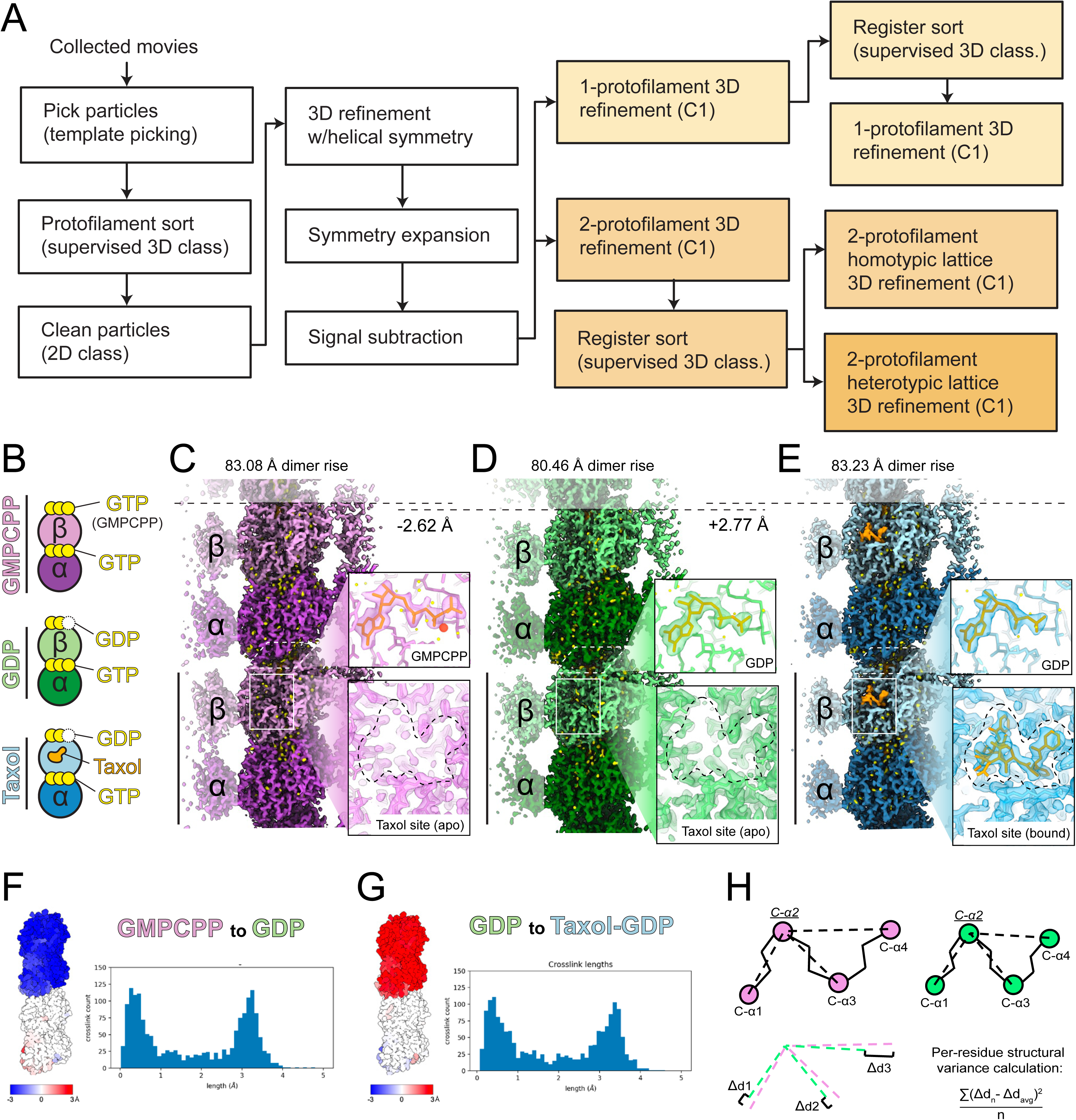
Overview of cryo-EM pipeline and 1-protofilament reconstructions. (A) Simplified overview of cryo-EM data processing pipeline. Light orange color steps are for 1-protofilament, orange for 2-protofilament homotypic lattices, and dark orange for 2-protofilament heterotypic lattices. (B) Compositional schematic of tubulin in each experimental condition. (C-E) Cryo-EM density of the GMPCPP (purple), GDP (green), and Taxol-GDP (blue) 1-protofilament reconstructions. Zoom-ins depict exchangeable-site nucleotides and Taxol-binding site. Ligands (nucleotides and Taxol) are orange, magnesium is red, and water molecules are yellow. (F & G) (left) 1-protofilament model colored by longitudinal displacement in the listed comparisons, where blue is downward displacement, red is upward displacement, and white is no displacement. (right) Histograms of total rmsd. (H) Visual depiction of per-residue structural variance. In this example, the value for C-α2 is being calculated.

**Extended Data Fig. 2.**
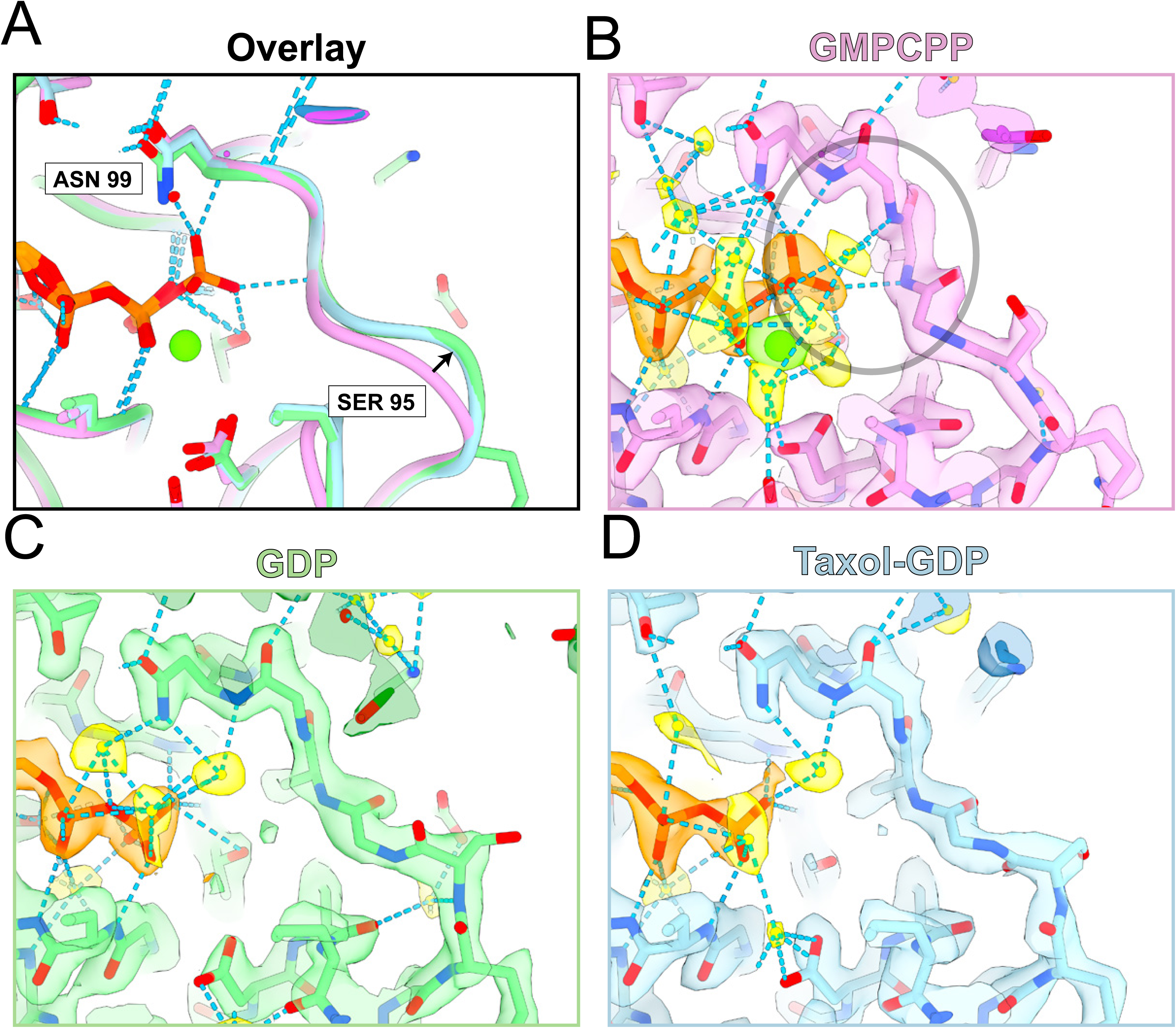
β-tubulin T3-loop conformation is nucleotide-dependent. (A) Structural overlay of the β-tubulin T3 ribbon model structures of GMPCPP (purple), GDP (green), and Taxol-GDP (blue). (B-D) Density and atomic representation of all structures. Nucleotides are orange, water molecules are yellow, magnesium ions are lime green, and hydrogen bonds are sky blue. Gray circle indicates terminal-phosphate-specific interactions with the GMPCPP T3 loop.

**Extended Data Fig. 3.**
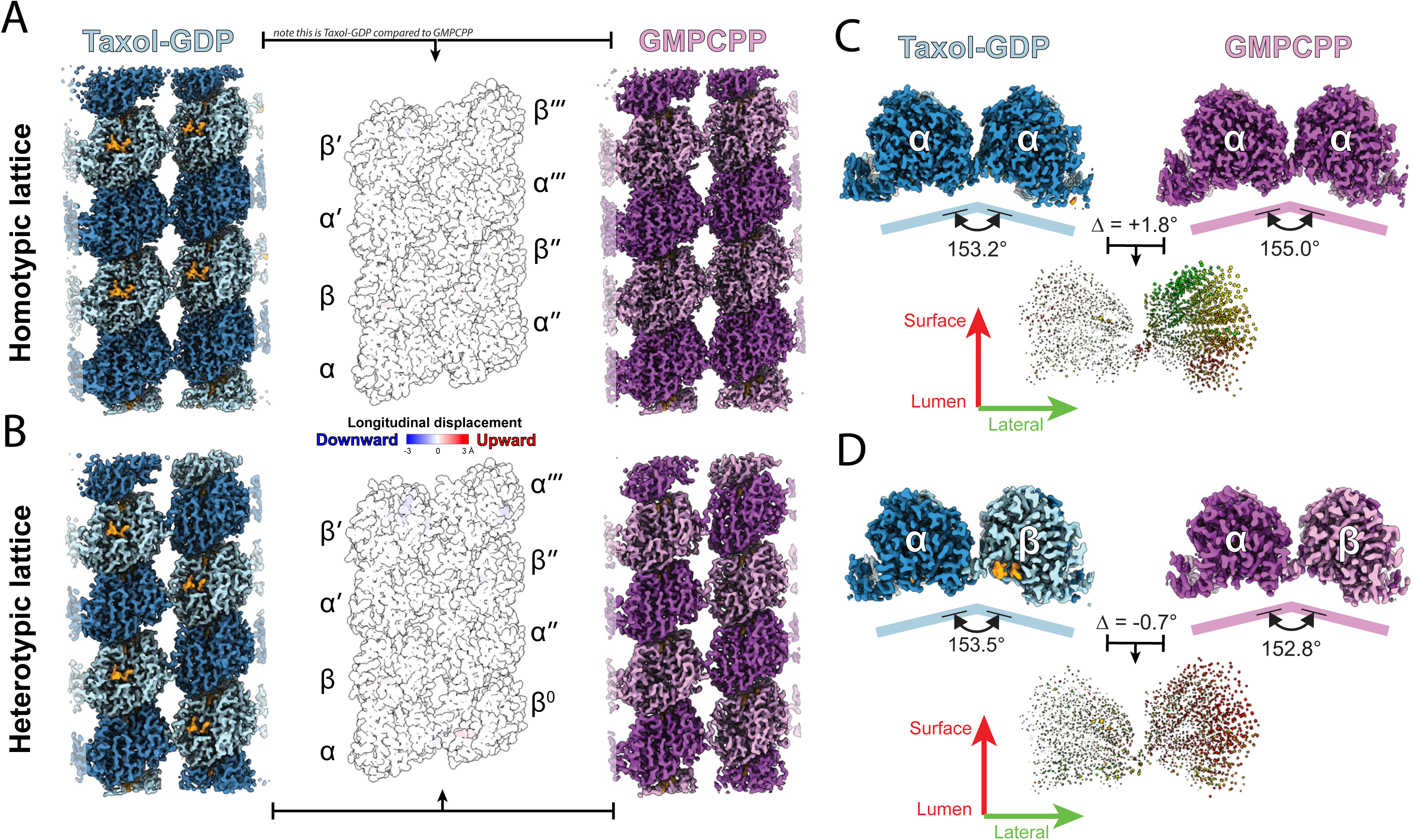
GMPCPP and Taxol-GDP 13-protofilament microtubules share lattice structures. (A & B) Longitudinal comparisons of GMPCPP and Taxol-GDP 2-protofilament homotypic (A) and heterotypic (B) lattices. 2-protofilament models colored by longitudinal displacement in the listed comparisons, where red is downward displacement, blue is upward displacement, and white is no displacement. (C & D) Lateral comparisons of GMPCPP and Taxol-GDP 2-protofilament homotypic (C) and heterotypic (D) lattices. (Bottom) Pairwise comparisons showing vectors between Cα positions. The vectors are dilated by a factor of distance/(max_distance*2) to emphasize larger relative changes and are colored according to the X and Y displacement with green and red, respectively.

**Extended Data Fig. 4.**
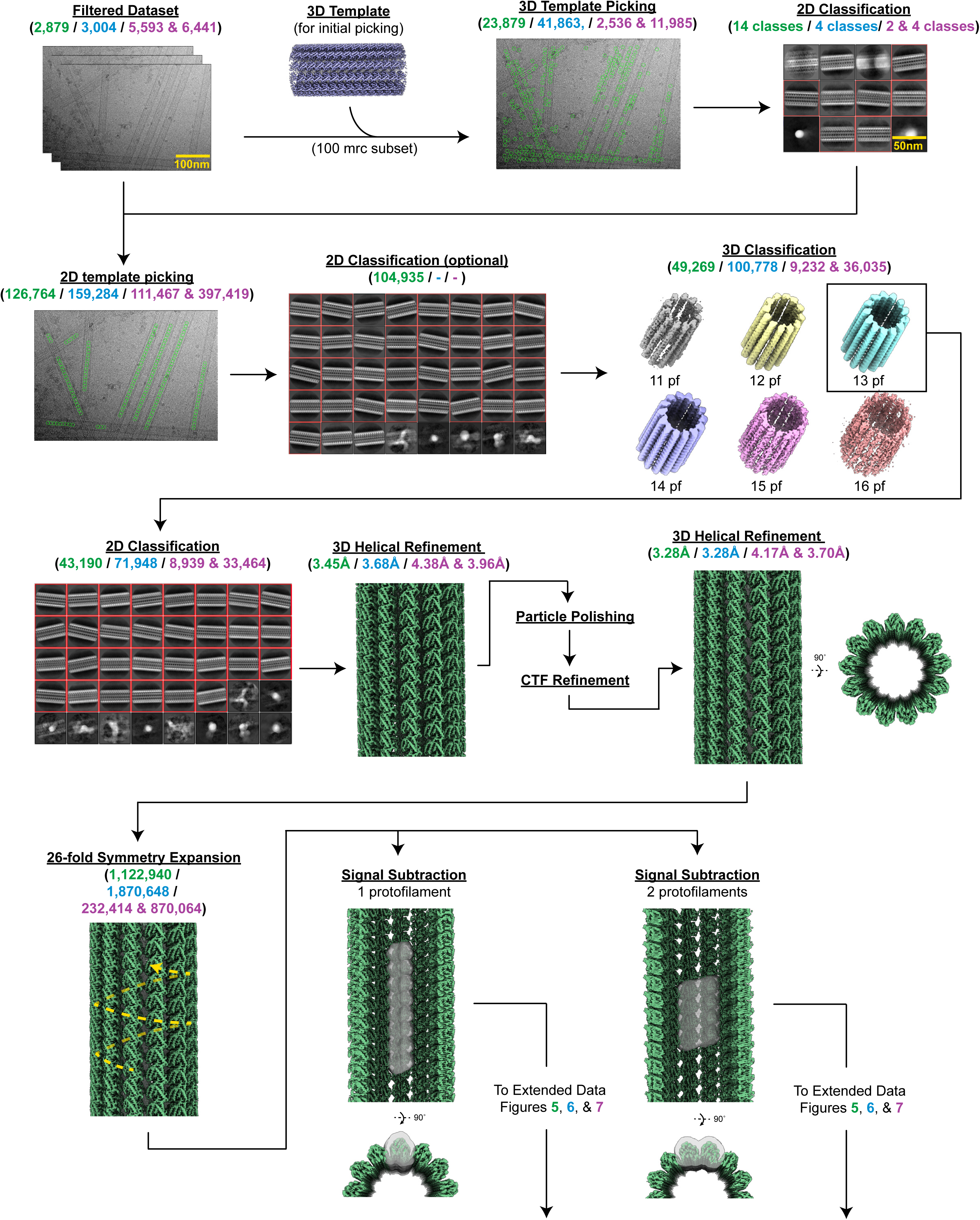
Whole-microtubule cryo-EM data processing workflow. Colored numbers in parentheses indicate particle number, class number, or resolution for a given experimental condition: green is GDP, blue is Taxol-GDP, and purple is GMPCPP (2 datasets).

**Extended Data Fig. 5.**
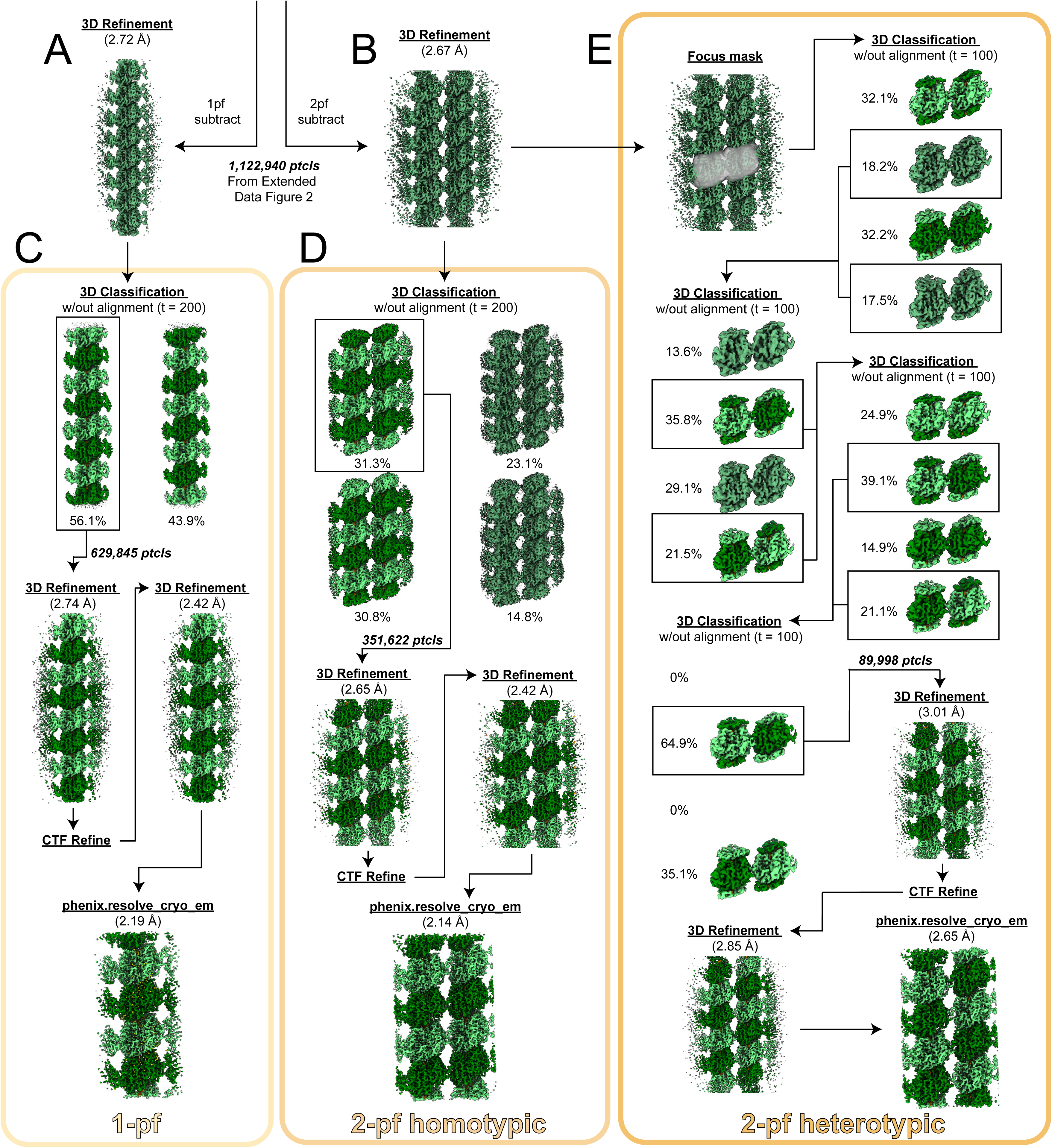
13-protofilament GDP microtubule cryo-EM data processing workflow. (continues from Extended Data Fig. 4) Cryo-EM processing workflows for 13-protofilament GDP microtubules, split into 1-protofilament (left, light orange), 2-protofilament homotypic (middle, orange), and 2-protofilament heterotypic (right, dark orange) lattice structures.

**Extended Data Fig. 6.**
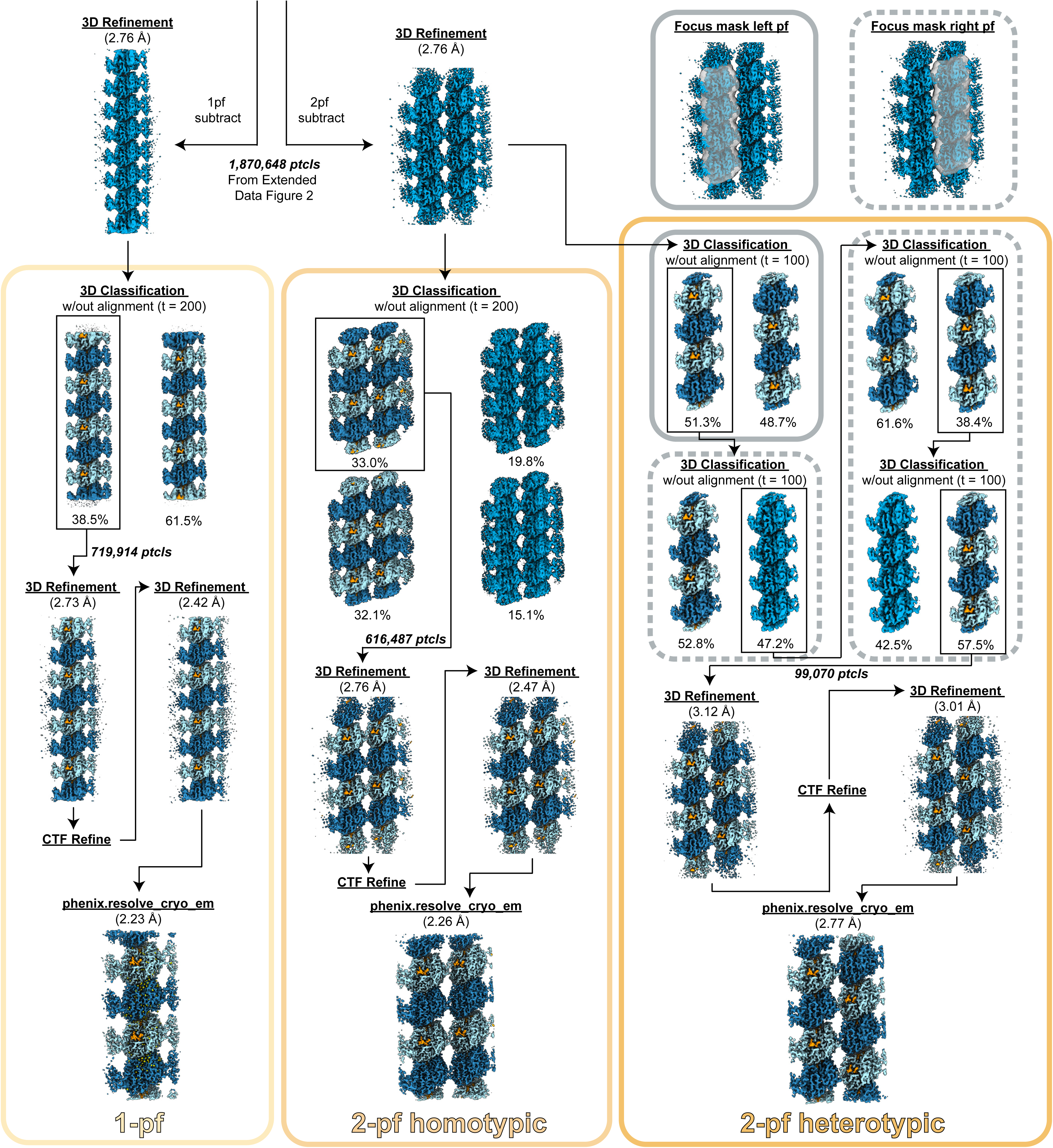
13-protofilament Taxol-GDP microtubule cryo-EM data processing workflow. (continues from Extended Data Fig. 4) Cryo-EM processing workflows for 13-protofilament Taxol-GDP microtubules, split into 1-protofilament (left, light orange), 2-protofilament homotypic (middle, orange), and 2-protofilament heterotypic (right, dark orange) lattice structures.

**Extended Data Fig. 7.**
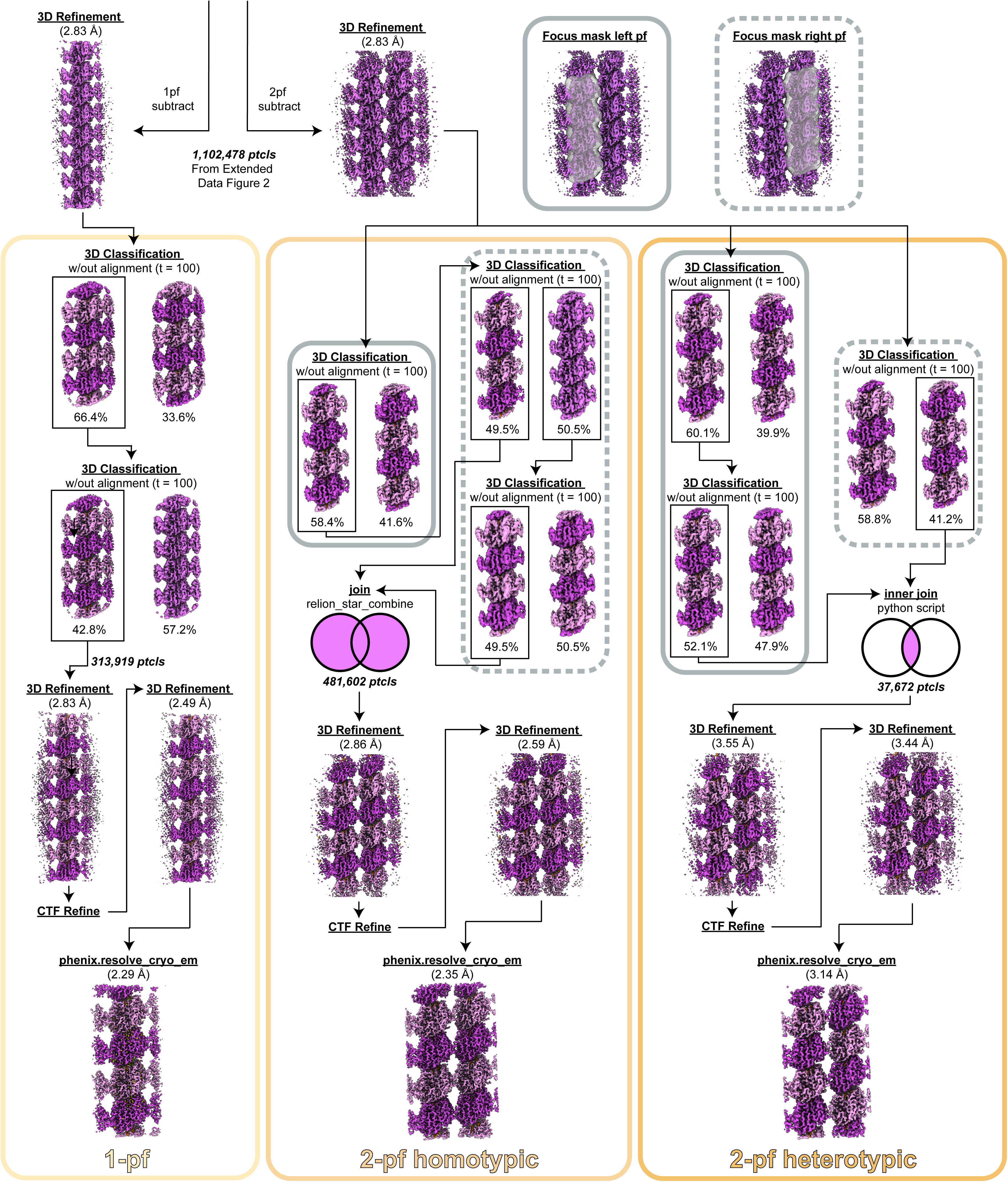
13-protofilament GMPCPP microtubule cryo-EM data processing workflow. (continues from Extended Data Fig. 4) Cryo-EM processing workflows for 13-protofilament GMPCPP microtubules, split into 1-protofilament (left, light orange), 2-protofilament homotypic (middle, orange), and 2-protofilament heterotypic (right, dark orange) lattice structures.

**Extended Data Fig. 8.**
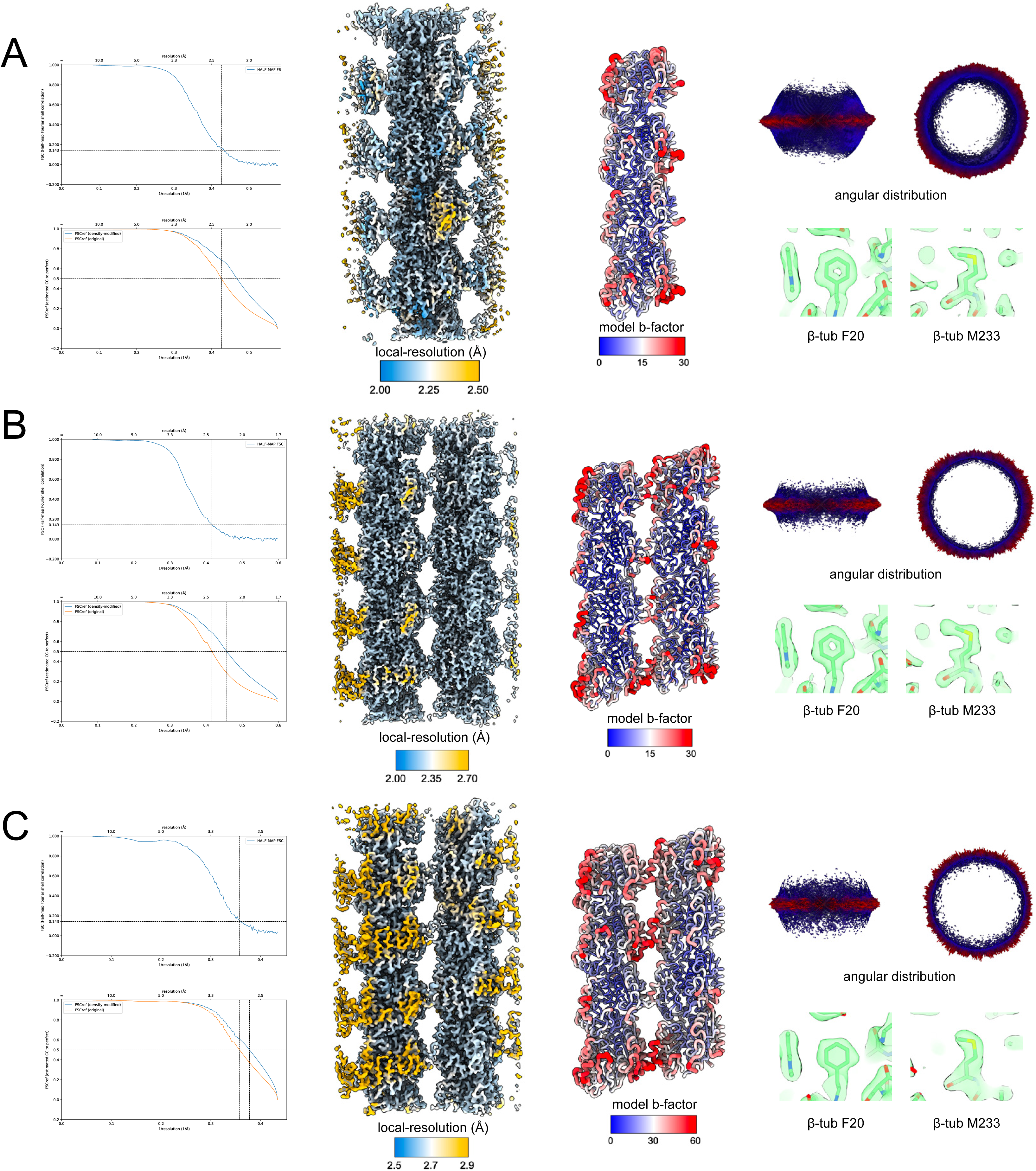
13-protofilament GDP microtubule structure validation. FSC curves for unsharpened (top) and density-modified (bottom) maps, final density map colored by local resolution, fitted model colored by b-factor, 3D particle orientation distribution, and zoom-in density comparisons for 1-protofilament (A), 2-protofilament homotypic (B), and 2-protofilament heterotypic (C) lattices structures.

**Extended Data Fig. 9.**
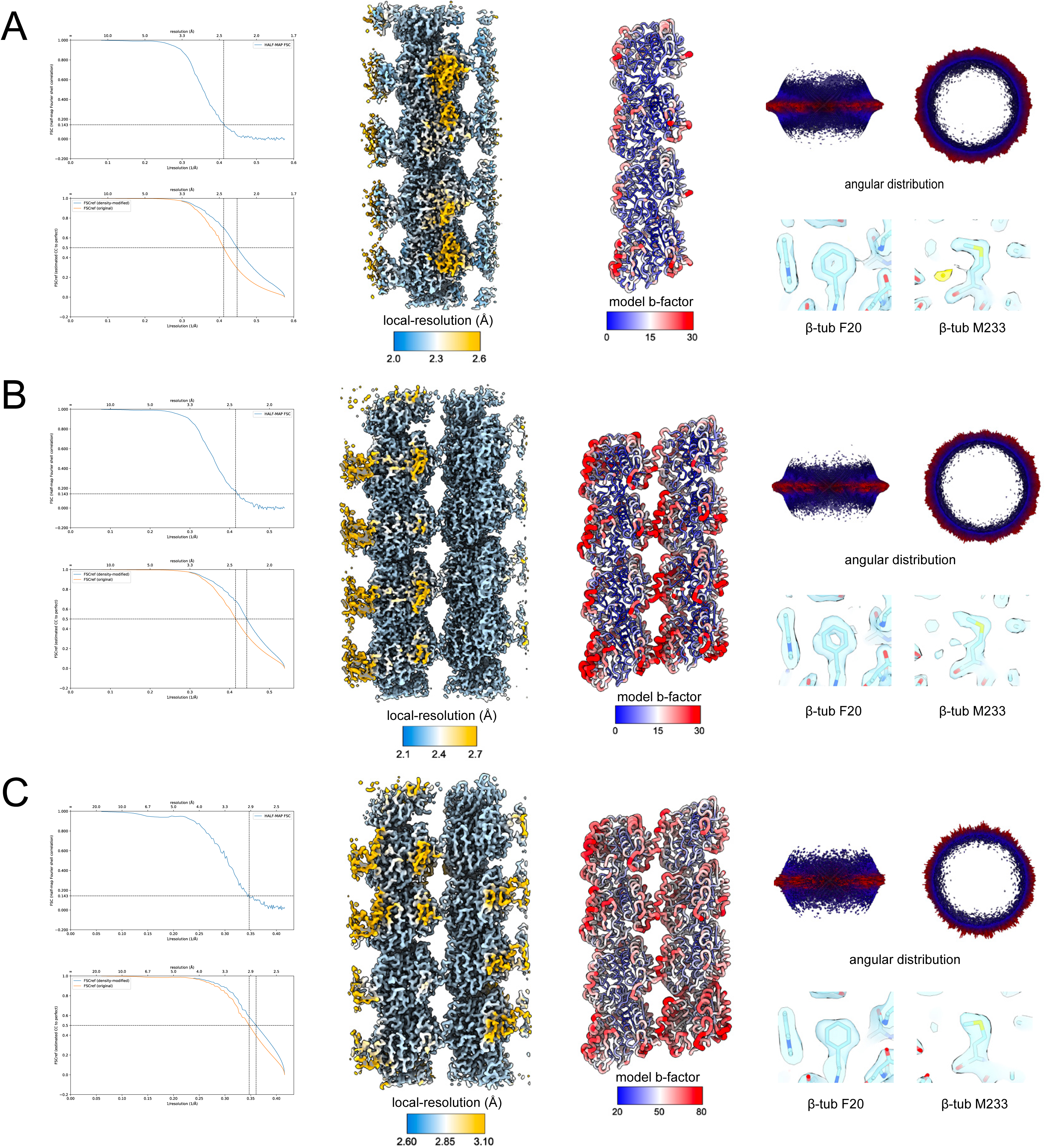
13-protofilament Taxol-GDP structure validation. FSC curves for unsharpened (top) and density-modified (bottom) maps, final density map colored by local resolution, fitted model colored by b-factor, 3D particle orientation distribution, and zoom-in density comparisons for 1-protofilament (A), 2-protofilament homotypic (B), and 2-protofilament heterotypic (C) lattices structures.

**Extended Data Fig. 10.**
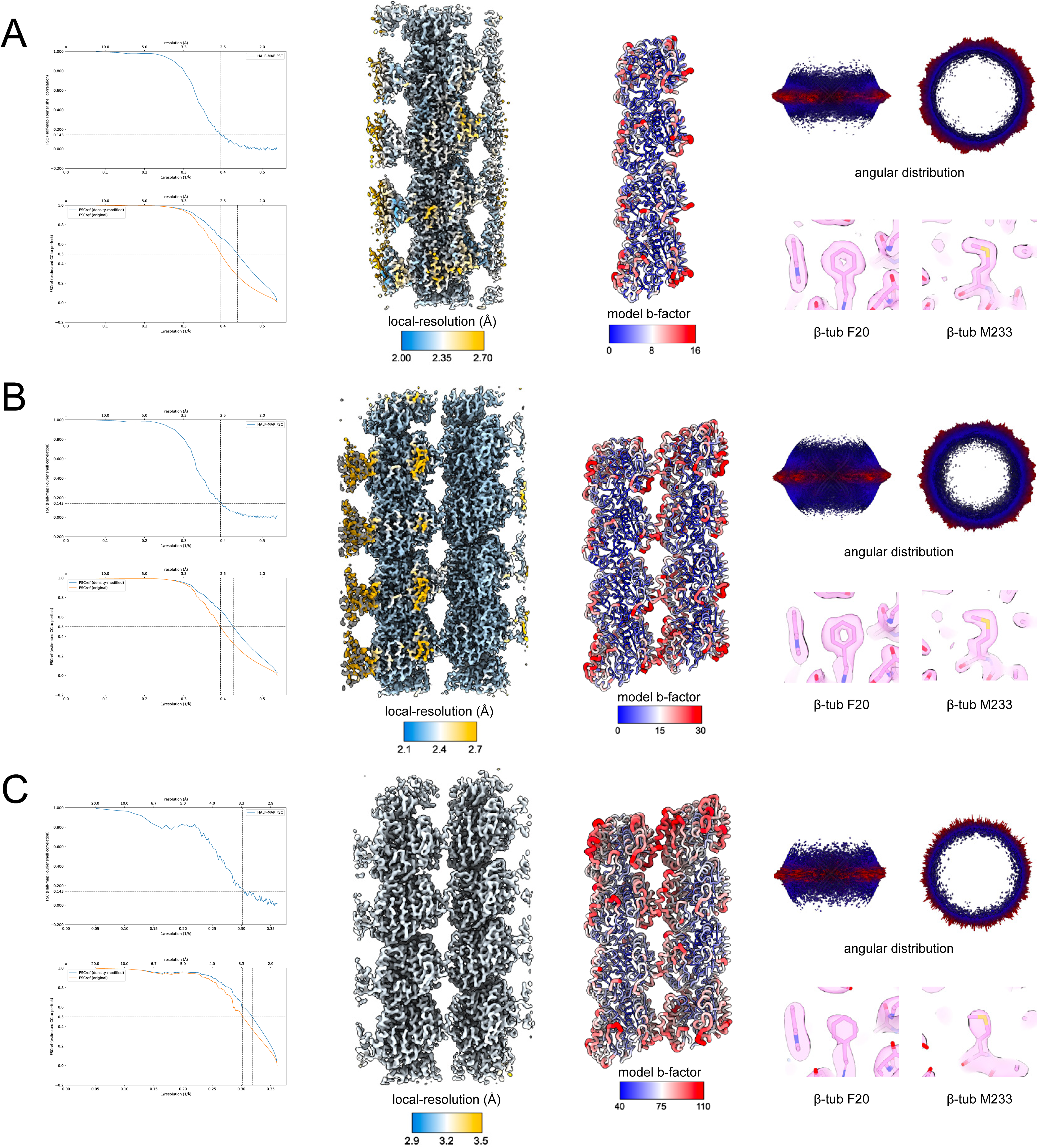
13-protofilament GMPCPP structure validation. FSC curves for unsharpened (top) and density-modified (bottom) maps, final density map colored by local resolution, fitted model colored by b-factor, 3D particle orientation distribution, and zoom-in density comparisons for 1-protofilament (A), 2-protofilament homotypic (B), and 2-protofilament heterotypic (C) lattices structures.

**Movie 1 - Hydrolysis switch morph.**

**Movie 2 - Compaction switch morph.**

**Movie 3 - Surface switch morph.**

**Movie 4 - Helix 7 - top morph.**

**Movie 5 - Helix 7 - bottom morph.**

**Movie 6 - Homotypic lattice morph - top view.**

**Movie 7 - Homotypic lattice morph - side view.**

**Movie 8 - Heterotypic lattice morph - top view.**

**Movie 9 - Heterotypic lattice morph - side view.**

